# Neural ensembles that encode affective mechanical and heat pain in mouse spinal cord

**DOI:** 10.1101/2024.02.05.578816

**Authors:** Ming-Dong Zhang, Jussi Kupari, Jie Su, Yizhou Hu, Kajsa A. Magnusson, Laura Calvo-En-rique, Dmitry Usoskin, Gioele W Albisetti, Andrew D Leavitt, Hanns Ulrich Zeilhofer, Tomas Hökfelt, Malin C. Lagerström, Patrik Ernfors

**Affiliations:** Department of Medical Biochemistry and Biophysics, Division of Molecular Neurobiology, Karolinska Institutet, Stockholm, Sweden; Department of Immunology, Genetics and Pathology, Uppsala University, Uppsala, Sweden; Institute of Pharmacology and Toxicology, University of Zurich, and Swiss Institute of Technology (ETH) Zurich, Zurich, Switzerland; Department of Medicine, University of California, San Francisco, CA, USA; Department of Laboratory Medicine, University of California, San Francisco, CA, USA; Department of Neuroscience, Karolinska Institutet, Stockholm, Sweden

## Abstract

Acute pain is an unpleasant experience caused by noxious stimuli. How the spinal neural circuits attribute differences in quality of noxious information remains unknown. By means of genetic capturing, activity manipulation and single cell RNA sequencing, we identified distinct neural ensembles in mouse spinal cord encoding mechanical and heat pain. Re-activation or silencing of these ensembles potentiated or stopped, respectively, affective but not reflex behaviour without altering pain behaviour to cross stimuli modality. Within ensembles, polymodal Gal^+^ inhibitory neurons with monosynaptic contacts to A-fiber sensory neurons gated affective pain independent of modality. Peripheral nerve injury led to microglia driven inflammation and an ensemble transition with decreased recruitment of Gal^+^ inhibitory neurons and increased excitatory drive. However, activating Gal^+^ neurons reversed hypersensitivity associated with neuropathy. Our results reveal the existence of a spinal representation which forms the neural basis of the discriminative and affective qualities of acute pain and that these neurons are under the control of a shared feed-forward inhibition.

## Introduction

Detection and response to intense mechanical force and heat enables a protection from what can cause injury^1-3^. Protective reflexes and the unpleasant experience of pain are initiated by the activation of specialized primary afferent sensory neurons with peripheral termini in the skin and central termini in the spinal cord. Within the spinal cord, the incoming information is processed before it is relayed to higher brain regions. A local neuronal network (i.e. ensemble) contacted by nociceptor terminals consist of excitatory and inhibitory interneurons as well as projection neurons. How spinal cord neurons transform nociceptive information into a neural computation representing the discriminative and affective qualities of pain remains unclear^4^. A better understanding of the neural principles of pain is essential for developing new therapeutic strategies to limit suffering in chronic pain patients. Several models have been proposed for coding nociceptive information in primary afferent sensory neurons, including dedicated molecular neuron types, necessity of temporal summation or patterns of firing, as well as various combinations of these ideas^4-7^. While in activity recordings of spinal projection neurons most display a polymodal response, some neurons respond only to noxious mechanical stimuli or heat^8-11^, illustrating the existence of a neural representation in the spinal cord conveying cutaneous mechanical and heat sensations. However, experimental studies with functional manipulation of populations of excitatory spinal neurons defined by their expression of neurochemical markers has revealed these to be polymodal, and thus affect both mechanical and thermal nociception or with marginal preference for coding modality-selective noxious stimuli^4,12-19^.

Embedded within these circuits, inhibitory neurons play a critical role^20^, as was originally proposed in the gate control theory of pain^21^ and a reduction of inhibitory activity induces allodynia and hyperalgesia across all sensory modalities and pain states similar to those associated with chronic pain^22^. Such disinhibition increases excitability of *Tacr1* expressing lamina I projection neurons known to transmit noxious information from the spinal cord to the brain^1,23,24^. Thus, spinal inhibitory neurons maintain appropriate activity levels in the neuronal circuits by regulating afferent sensory information on the way to the output projection neurons. Recently, populations of spinal cord inhibitory neurons have been functionally accessed based on marker gene expression. Among those expressed in adult mice are dynorphin and parvalbumin expressing neurons, which prevent touch inputs from activating pain circuits in the superficial dorsal horn. Consistently, preventing the activity of these neurons leads to a marked reduction in the threshold for the mechanically evoked withdrawal reflex^4,14,25^. However, little is known about inhibitory mechanisms involved in affective qualities of pain.

Activity-dependent expression of the Fos gene in the spinal cord marks neural activity caused by peripheral noxious stimuli^26,27^. Although, efforts have been made to decode those activated neurons in spinal cord using different genetic strategies, there has been limited progress^16,28,29^. Single cell RNA-sequencing (scRNA-seq) has revealed a marked transcriptional heterogeneity including up to 30 excitatory and inhibitory neuron types in the dorsal horn of the spinal cord^30-32^. A molecular atlas of spinal neurons and genetic strategies that captures active neurons with high temporal resolution open for functional and molecular studies that can provide insights into the neural substrates that represent noxious information in the spinal cord. Here, we identify the neural ensembles in the spinal cord coding for the affective dimension of noxious mechanical and heat stimuli.

## Results

### Noxious mechanical and heat stimuli encoded in spinal ensembles

We first performed fluorescent in situ hybridization and immunohistochemical studies for *Fos* gene expression to establish robust and reproducible stimulation paradigms for activation of spinal neurons by peripheral mechanical or heat stimuli. With these used stimulation protocols, *Fos*^+^ (mRNA) and c-Fos^+^ (protein) neurons were systematically found in the medial superficial layers of the corresponding lumbar spinal dorsal horn receiving inputs from the hind paw (ExtendedData Fig. 1a and Fig. 1a). To identify and access those mechanically or heat active neurons for manipulation, we used the stimulus-coupled transcription technology CANE (capturing activated neuronal ensembles) which affords very high temporal resolution^33^. Specificity of the Fos^dsTVA^ mouse strain used in the CANE technology was validated by determining co-localization of injury induced c-Fos and TVA (tumor virus A) expression in superficial layers of spinal cord and brain (dentate gyrus and supraoptic nucleus) and revealed a very high fidelity by means of TVA and c-Fos co-localization cells (Fig. 1b and ExtendedData Fig. 1b). Fos^dsTVA^ mice bred to ROSA26^Tomato^ reporter mice (Fos^dsTVA*^R26^Tom^) were used to ascertain the accuracy of genetically captured active neurons in the spinal cord. Mice (Fos^dsTVA*^R26^Tom^) were administered mechanical or heat stimuli and intra-spinally injected of a Cre expressing EnvA pseudotyped lentivirus (EnvA^M21^ lenti-Cre virus), thus limited transduction only to the cells that transiently express TVA^34^. Three weeks later the same mechanical and heat stimuli were administered again, and animals sacrificed for analysis of expression of Cre-dependent Tomato reporter initiated at the first stimulations with c-Fos induced by the second stimulations (Fig. 1c, d). The overlaps of active ensembles captured by EnvA^M21^ lenti-Cre virus (“TRAPed” neurons labelled with Tom) for the first stimulus and c-Fos for the second stimulus were 51.78 ± 4.69% and 68.59 ± 4.54% for mechanical and heat, respectively (Fig. 1d).

**Fig. 1:**
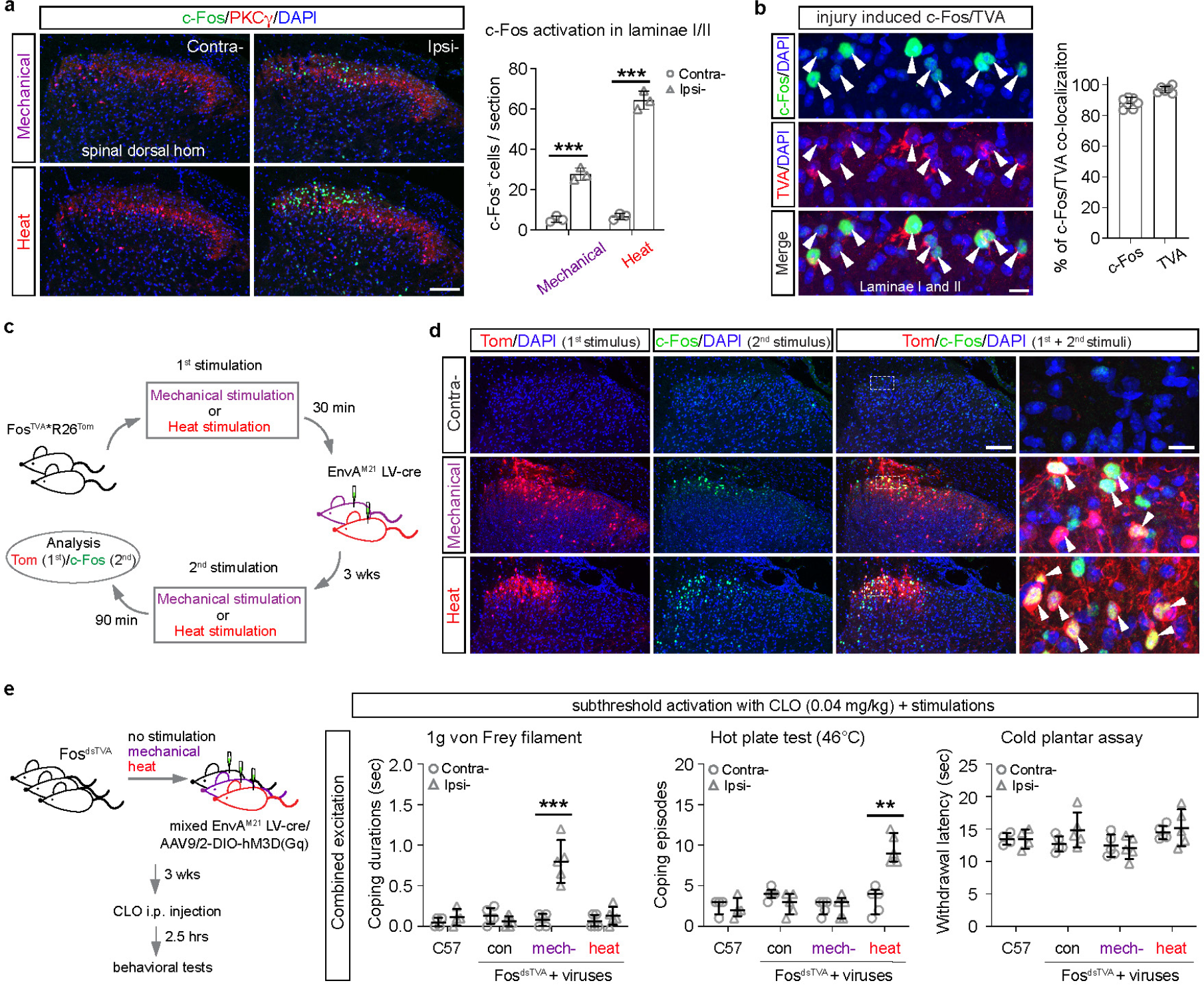
Capturing ensembles coding for mechanical and heat pain in spinal dorsal horn. **a**, Representative images of c-Fos expression with immunohistochemistry in medial spinal dorsal horn under noxious mechanical or heat stimulus applied to the left hind paw of wild type mice. PKCγ was used as an anatomical land maker, which specifically labels a group of excitatory interneurons mainly in the inner layer of lamina II (IIi) of spinal cord. Number of c-Fos positive neurons from laminae I and II per section under each stimulation condition was quantified (n = 6 mice). **b**, Colocalization of destabilized TVA (dsTVA) and c-Fos in spinal cord of FosdsTVA^*^R26Tom mice induced by skin incision and/or nerve injury. The proportion of co-localized neurons to Fos or TVA neurons were quantified (n= 6 mice). **c**, Schematic representation of the working flow of validation experiment for the same type of stimulus, where the ensembles from 1st stimulation was labeled with Tomato through EnvAM21 lenti-cre virus and the ensembles from 2nd stimulus was labeled with c-Fos. **d**, Representative images from immunohistochemistry show ensembles activated and labeled with Tomato (Tom) and c-Fos in spinal dorsal horn from the same stimuli as illustrated in (c), respectively (n = 6 mice). Contralateral (Contra-) spinal cord from both heat and mechanical stimulation experiments showed similar results, where c-Fos expression was barely detected. High magnification images from the inset box show the co-localization of Tom and c-Fos indicated by arrowheads. **e**, Left, schematic diagram for the functional study of gain of function for ensembles infected by AAV9/2-DIO-hM3D(Gq) through subthreshold clozapine (CLO) combined with peripheral applied stimuli. DIO: double-floxed inverted open-reading frame. Right, coping durations/episodes for 1g von Frey filament, hot plate test (46°C) or reflex latency for cold plantar assay under subthreshold activation of mechanical or heat ensembles (n = 19 mice). Con: FosdsTVA mice injected virus without stimulation; C57: wild type mice; Contra-: contralateral; Ipsi-: ipsilateral. Scale bars: 100 μm in (**a**) and (d), 25 μm in (**b**) and inset in (**d**). Data are expressed as mean ± standard deviation. Differences between contra- and ipsi- from different groups were analyzed by ordinary one-way ANOVA followed by Bonferroni’s multiple comparisons test (**a, e**). The coping episodes for Hot plate test in (**e**) were presented as median with interquartile range and differences between contra- and ipsi-from each group were analyzed by Mann-Whitney test (two-tailed). ^***^ indicates p < 0.001 and ^**^ indicates p < 0.01.

We next examined the sufficiency of the mechanical and heat ensembles for pain behaviors and whether sub-threshold activation resulted in cross-modality effects using Fos^dsTVA^ mice intraspinally injected with a virus mixture of EnvA^M21^ lenti-Cre and AAV9/2-hEF1a-DIO-hM3D(Gq)-mCherry. When such mixed viruses were intraspinally administered to these mice co-incident with the mechanical or heat stimuli, it led to activity-dependent, temporally, and spatially controlled DNA recombination and hM3D(Gq) DREADD expression. A subthreshold dose of clozapine (CLO) which resulted in no/rare coping behaviours (paw shaking, lifting/guarding or licking) by itself in wild-type control (C57/Bl6), control Fos^dsTVA^ mice (no stimulation but with virus injected) or Fos^dsTVA^ mice (stimulation and virus injected) was titrated. A dosage of 0.04 mg/kg clozapine through intraperitoneal injection (2.5 hr - 4.0 hr post injection) was chosen as the subthreshold dose for coping behaviors. When subthreshold activation was combined with natural stimuli which normally resulted in little coping behaviour, a robust coping was observed within modality and without any effects across modalities, indicating these ensembles to code also for intensity (Fig. 1e). Interestingly, subthreshold activation also resulted in moderate effects on reflex behaviour (ExtendedData Fig. 1c), suggesting that forced activation can weakly access reflex protective pathway.

To further establish the causal role of the mechanical and heat ensembles for pain behaviors, we used the inhibitory hM4Di-DREADD, whose expression is Cre-dependent in the Fos^dsTVA*^R26^PDi^ mouse strain. Thus, using such mice we tested the requirement of the ensembles for pain behavioural responses and determined whether silencing the neurons activated by one noxious stimulus altered responses to other types of noxious stimuli.

We first genetically captured and expressed hM4Di-DREADD in the spinal ensemble activated by noxious mechanical stimuli. Silencing the ensemble three weeks later by the agonist clozapine led to a pronounced loss of coping behaviour (including shaking, lifting and licking) to pricking pain (2g von Frey filament) with a temporal effect corresponding to peak plasma concentration and clearance of clozapine. In contrast, coping behaviour to noxious heat (50°C hot plate) or cold (pre-cold acetone) was unchanged (Fig. 2a and ExtendedData Fig. 2a). While coping behaviour is believed to soothe suffering invoked by pain, reflexive defensive behaviour is a protective response that prevent or limit injury^17^. Mice in which the noxious mechanical ensemble was silenced showed no changes in withdrawal latency to heat, cold or thresholds to punctate mechanical force evoked by von Frey filaments (ExtendedData Fig. 2a). Thus, this mechanical ensemble of spinal neurons transforms afferent no-ciception information into a signal necessary for affective pain behaviour to noxious mechanical stimuli.

**Fig. 2:**
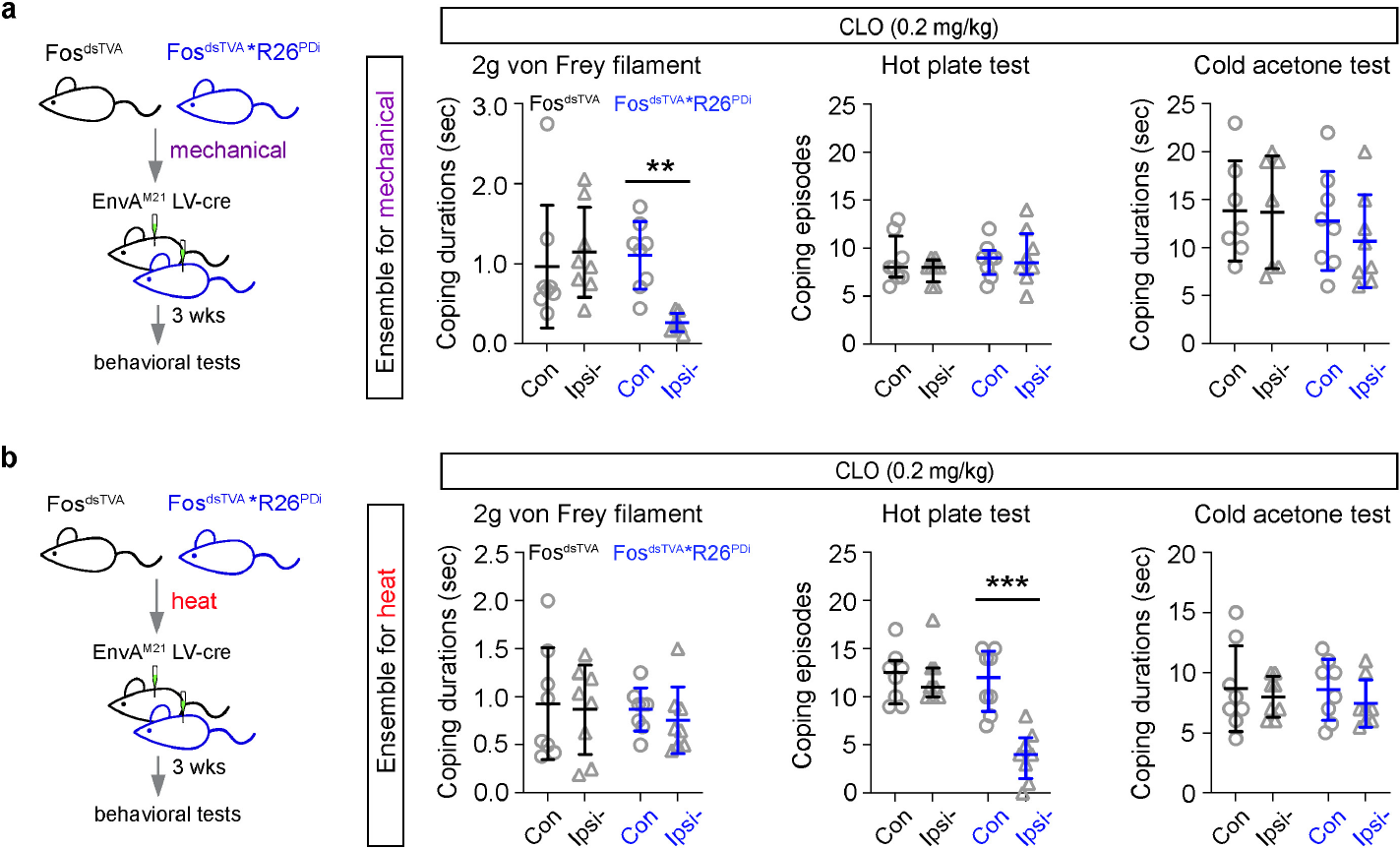
Chemogenetic inhibition of ensembles coding for mechanical and heat pain in spinal cord. Schematic diagram for chemogenetic inhibition of ensembles that code for mechanical pain (**a**) and heat pain (**b**) in spinal dorsal horn with intraspinal injected pseudotyped EnvA^M21^lenti-cre virus in Fos^dsTVA*^R26^PDi^ and Fos^dsTVA^ (control) mice (n = 32 mice, 8 mice in each group). Coping behavior (durations or episodes) was assessed with 2g von Frey filament test (mechanical), hot plate test (heat, 50°C) and cold acetone test (cold) after 60 min of intraperitoneal administration of clozapine (CLO, 0.2 mg/kg) and summarized in (**a**) and (**b**). Fos^dsTVA^ mice were used as a control group. Con: contralateral, Ipsi-: ipsilateral. Data are expressed as mean ± standard deviation coping durations and median with interquartile range for coping episodes. Coping duration differences between contra- and ipsi-from different groups were analyzed by ordinary one-way ANOVA followed by Bonferroni’s multiple comparisons test (**a, b**). The coping episodes differences between contra- and ipsi-from each group were analyzed by Mann-Whitney test (two-tailed).^***^ indicates *p* < 0.001 and ^**^ indicates *p* < 0.01.

We next used the same capturing strategy to express hM4Di in spinal neurons activated by noxious heat. Silencing the heat ensemble three weeks later by the agonist clozapine did not lead to any changes in coping behavior in response to noxious pricking- or cold-induced pain while a near complete absence of noxious heat pain behaviour was observed, with an effect that corresponded to peak plasma concentration and clearance of clozapine (Fig. 2b and ExtendedData Fig. 2b)^35^. There was no effect on reflexive defensive responses to heat, cold or withdrawal thresholds to punctate mechanical force evoked by von Frey filaments (Fig. 2b and ExtendedData Fig. 2b). Intraspinal injection of EnvA^M21^ lenti-Cre virus did not affect behavior in control Fos^dsTVA^ mice and clozapine alone had no effect on contralateral limb pain behavioural in any of the experiments (Fig. 2A, B). Similar results were obtained in ablating instead of silencing the heat ensemble for mechanical and heat pain in the Fos^dsTVA*^R26^DTA^ mice (ExtendedData Fig. 2c). This shows the existence of parallel ensembles in the spinal cord that are required for mechanical and heat pain without affecting reflex responses.

### Molecularly defined ensembles

To determine the neural basis for encoding mechanical and heat pain signals in spinal cord, we employed scRNA-seq to identify the noxious mechanical and heat ensembles. Previous single cell transcriptomic studies of mouse spinal dorsal horn used to identify the molecular diversity of neuron types are either hampered by relatively few whole sequenced cells from young animals, reduced quality due to nuclei sequencing or loss of variability due to an integration of whole cell and nuclei datasets across different RNA sequencing platforms^30-32,36-38^. Identification of the mechanical and heat ensembles depends on the quality of the spinal cord atlas and the age of animals. We performed whole cell sequencing of lumbar spinal dorsal horn neurons from adult BAF53b-Cre^*^R26^Tom^ mice (ExtendedData Fig. 3a, b), where all neurons are labelled with Tom^39^. The new adult spinal dorsal horn neuronal atlas contained 18,590 neurons (4,926 median genes detected) which clustered into 10 inhibitory (In1-9, In18) and 17 excitatory (Ex10-17, Ex19-Ex27) neuron types (Fig. 3a and ExtendedData Fig. 3c-f). This new spinal neuronal atlas provided a more powered dataset to explore cell types with a number of unique markers defining each neuron type (ExtendedData Fig. 4a), including small cell clusters e.g. *Pkd2l1*^+^ In1 (*Slc32a1*^+^;*Scl6a5*^-^, Cerebrospinal fluid-contacting neurons, ExtendedData Fig. 4a) in lamina X compared to previous annotations^30,32,40^. A machine learning classifier (scPred)^41^ was trained with about two thirds of the neurons and tested on the remaining annotated neurons with a prediction accuracy of 91.94% (ExtendedData Fig. 4b). This classifier was also used to establish the relationship between the current and Häring’s^30^ atlases and showed high prediction accuracy also across the datasets and most cells to be identified in both, with a few exceptions (ExtendedData Fig. 4c, d). As a result of increased sequencing depth (4,926 vs 3,387 genes) and numbers of cells (18,590 vs 1,545 cells), the current atlas showed a marked improvement in cell type resolution based on expression of excitatory/inhibitory marker genes, genes coding neuropeptides and selected marker genes for clusters (ExtendedData Fig. 5a-f, Supplementary Tables 1 and 2).

**Fig. 3:**
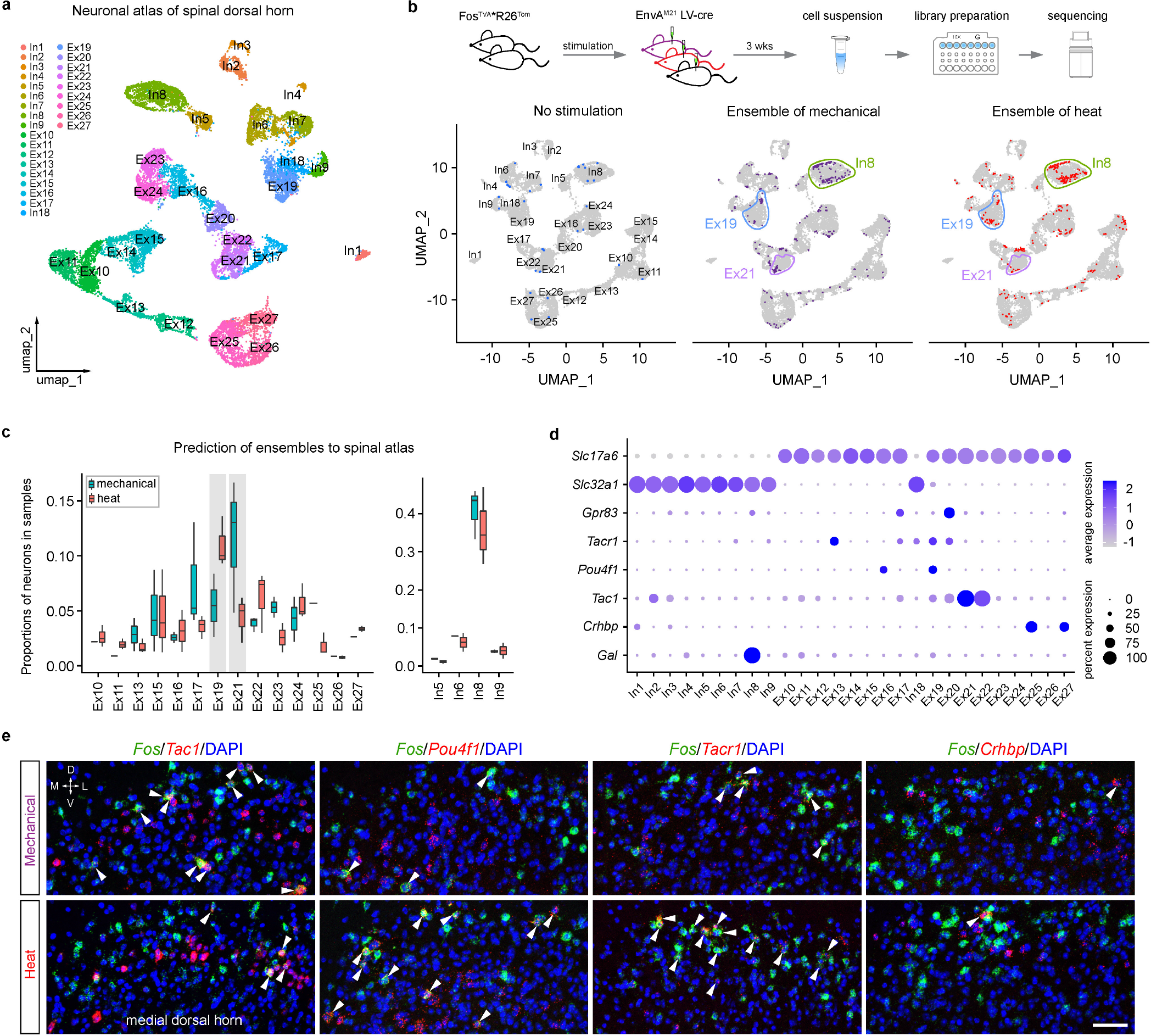
Decoding ensembles coding for mechanical and heat pain with single cell RNA-seq. **a**, UMAP plot composed of 18,590 single cell RNA-sequenced spinal dorsal horn neurons from adult mice (BAF53b-Cre^*^R26^Tom^, 12-20 weeks old, n = 8 mice) representing 27 clusters with 10 inhibitory (In1-9, In18) and 17 excitatory (Ex10-17, Ex19-27) cell types in adult spinal neuronal reference atlas. **b**, Top, schematic diagram for scRNA-seq of ensembles coding for mechanical and heat pain. Bottom, sequenced neurons from control, mechanical and heat stimulations were plotted on reference atlas by label transfer (n = 32 mice). Modality specific or shared cell types were highlighted with circles. **c**, Percentage of cell types from mechanical and heat ensembles were plotted for comparison. Shades of gray highlights modality specific cell types. **d**, Dot plot of expression of marker genes for excitatory neurons *Slc17a6* (vGLUT2), inhibitory neurons *Slc32a1* (VGAT), projection neurons *Gpr83* and *Tacr1*, and for specific cell types *Pou4f1*(Ex19/Ex16), *Tac1*(Ex21/Ex22), *Crhbp* (Ex25/27), *Gal* (In8). **e**, Representative images for mRNA co-expression of genes *Tac1, Pou4f1, Tacr1* or *Crhbp* with *Fos* in medial spinal dorsal horn from mechanical and heat stimulations. DAPI was used as counter staining. Arrowhead indicates co-localization. Scale bar = 50 μm. Data are expressed as mean ± s.d. in (**c**).

**Fig. 4:**
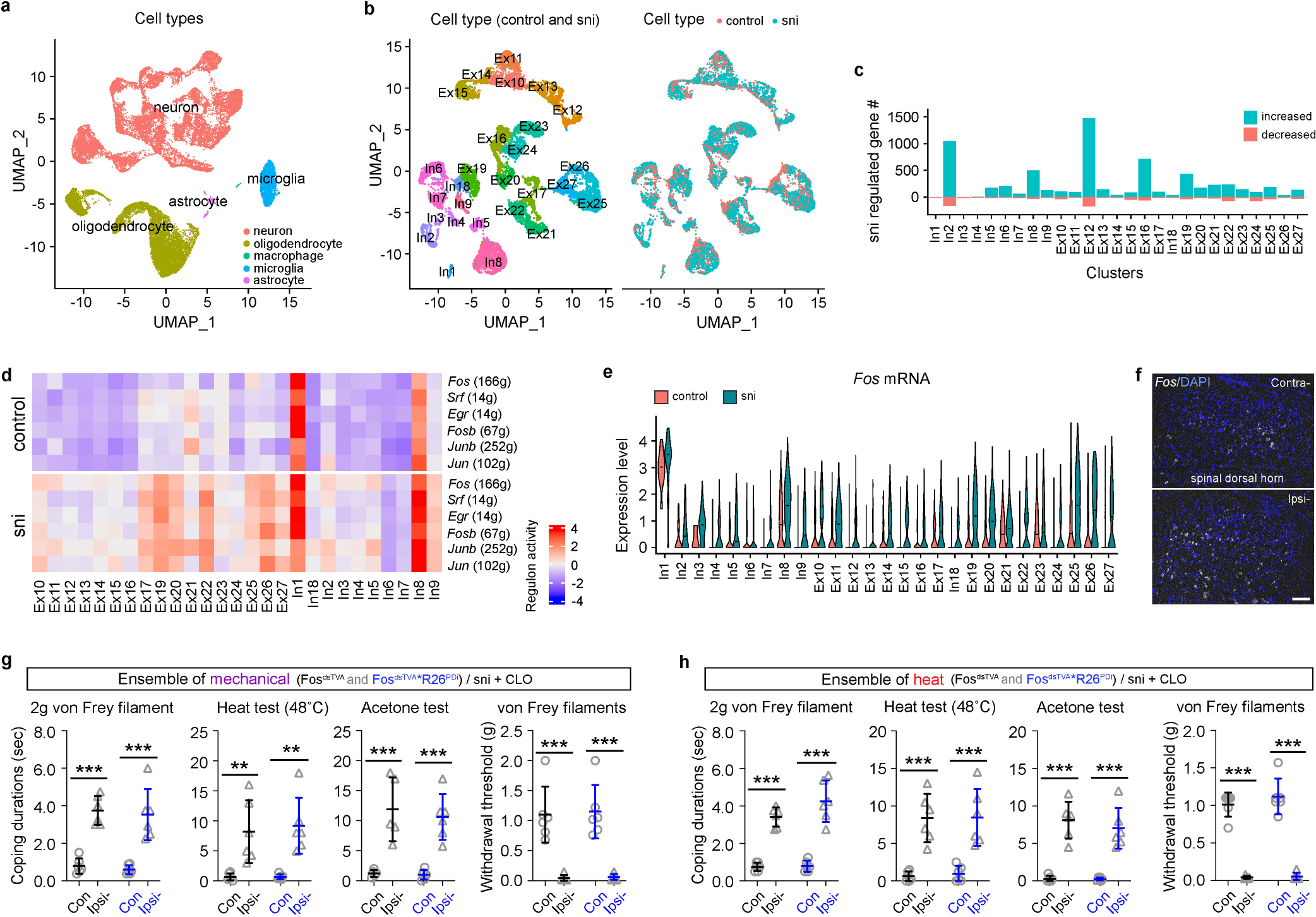
Spinal ensemble transition from nociceptive to chronic pain. **a**, UMAP plot is composed of 28,195 single cell RNA-sequenced cells (n = 4 mice for control and n = 4 for sni 6 weeks). **b**, UMAP plot shows neuronal cell types of 17,229 neurons (left) from both sni (11,234) and control (5,995) conditions (right). **c**, Numbers of regulated genes by sni (sni vs con) for each cell type by pseudobulk analysis are listed. Genes with Log_2_Fold-Change > 0.59 or < -1.0 and padj < 0.01 were counted as sni regulated. **d**, Heatmap of immediate early genes related regulons’ activities in neuronal cell types from control and sni from SCENIC analysis. **e**, Violin plot shows *Fos* expression in control and sni conditions through all neuronal cell types. **f**, Representative image of *Fos* mRNA in contralateral and ipsilateral spinal dorsal horn after sni (n = 4 mice). DAPI was used as counter staining. Scale bar = 100 μm. **g** and **h**, Coping durations for 2g von Frey filament, heat test, and acetone test or withdrawal threshold for von Frey filaments were recorded for effects of inhibition of mechanical and heat ensembles by clozapine (CLO) after sni (n = 24 mice, 6 mice/group). Data are expressed as mean ± standard deviation. Differences between contra- and ipsi- within the group were analyzed by ordinary one-way ANOVA followed by Bonferroni’s multiple comparisons test (**g, h**). ^***^ indicates *p* < 0.001 and ^**^ indicates *p* < 0.01.

**Fig. 5:**
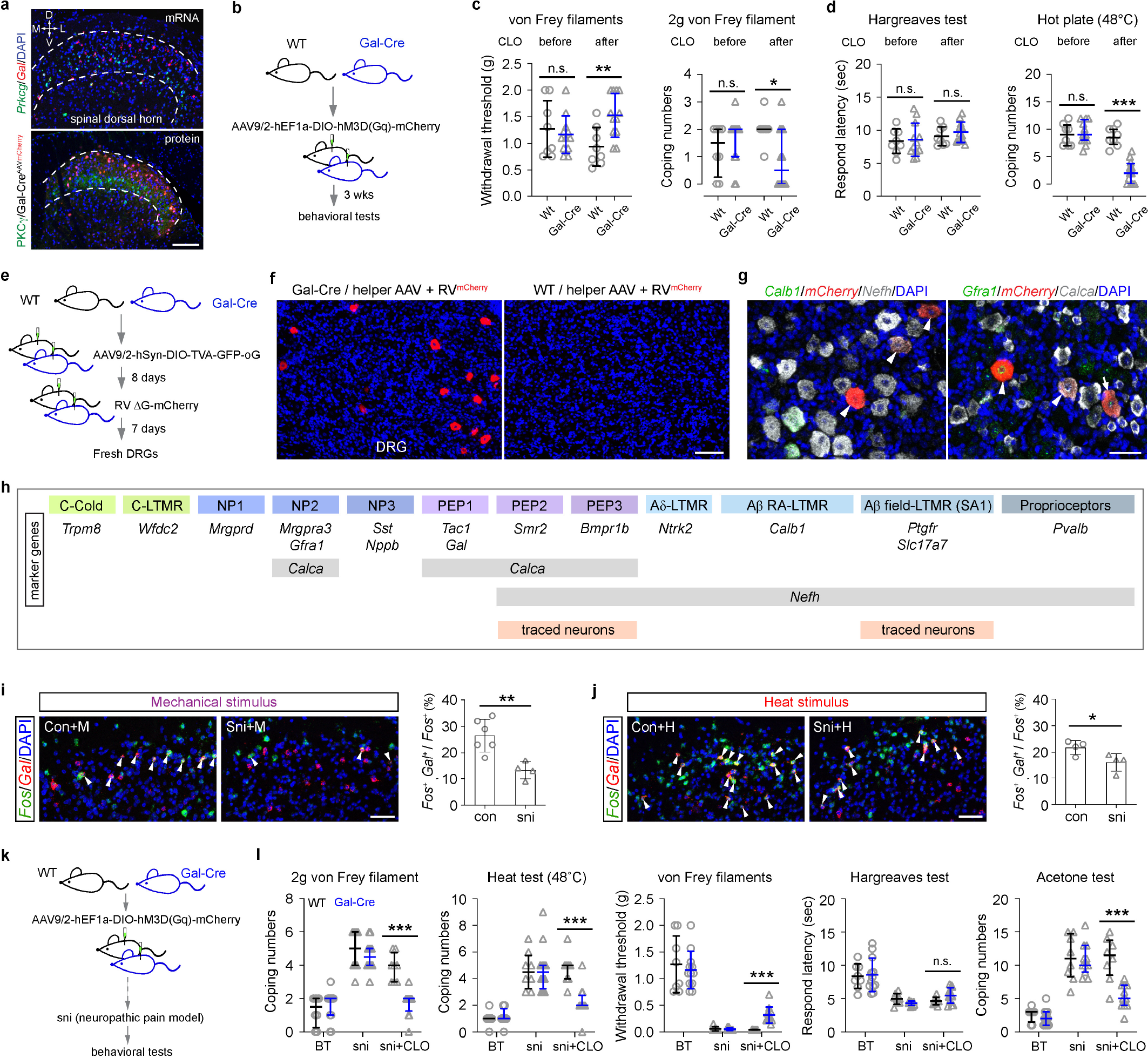
Gal+ In8 modulates both nociceptive and chronic pain. **a**, Top, endogenous expression of *Gal* and *Prkcg* mRNA in spinal dorsal horn of wt mice. Bottom, exogenous expression of mCherry through intra-spinal injected AAV9/2-DIO-mCherry in Gal-Cre mice, where PKCg was used to label the inner layer of lamina II (IIi) of spinal cord. **b**, Schematic diagram for gain of function for *Gal*^+^ In8 in behavioral tests through chemogenetics in (**c**) and (**d**). **c**, Withdrawal threshold of von Frey filaments and coping numbers of 2g von Frey filament was assessed before and after administration of clozapine (CLO) in wt and Gal-Cre mice injected with AAV virus. **d**, Response latency to Hargreaves test and coping numbers to hot plate test were assessed before and after administration of clozapine in wt and Gal-Cre mice injected with AAV virus. n= 8 mice for wt, n = 12 mice for Gal-Cre. **e**, Schematic diagram for monosynaptic retrograde tracing from *Gal*^+^ neurons in spinal dorsal horn to dorsal root ganglia (DRG) neurons with helper AAV virus and rabies virus. **f**, Retrograde traced DRG neurons were labelled with mCherry in Gal-Cre mice, where wt mice were used as control (n = 4 mice, 2 mice/group). **g**, Identification of traced mCherry^+^ DRG neurons with the combinations of RNAscope probes *Calb1*/*mCherry*/*Nefh* and *Gfra1*/*mCherry*/*Calca*. Arrows indicate triple labelling and arrowheads indicate double labelling. **h**, Summary for identified traced neurons in accordance with marker genes of DRG neuronal subtypes. **i**, Co-localization of *Fos*^+^ and *Gal*^+^ neurons in medial spinal dorsal horn after mechanical stimulation from control (n = 5) and sni (n = 4) mice. **j**, Co-localization of *Fos*^+^ and *Gal*^+^ neurons in medial spinal dorsal horn after heat stimulation from control (n = 4) and sni (n = 4) mice. Arrowheads indicate neurons with co-localization of *Fos* and *Gal*. **k**, Schematic diagram for effect of gain of function for *Gal*^+^ In8 neurons in sni induced chronic pain. **l**, Coping numbers for 2g von Frey filament, heat test, and acetone test or withdrawal threshold for von Frey filaments and response latency for Hargreaves test were recorded for effects of boost inhibition from *Gal*^+^ In8 neurons after sni. These sni mice were the same as in (**b**-**d**) and the basal threshold (BT) were the same values as those before clozapine for the same tests. DAPI was used as a counterstaining for nuclei in (**a, f, g, i** and **j**). Scale bars: 100 μm in (**a** and **f**), 50 μm in (**g, i** and **j**). Data are expressed as mean ± standard deviation, whereas coping numbers are expressed as median with interquartile range. Differences between before and after clozapine in (**c**) and (**d**) or before and after clozapine under sni condition between wt and Gal-Cre mice (**l**) were analyzed by ordinary one-way ANOVA followed by Bonferroni’s multiple comparisons test. The coping number differences were analyzed by Mann-Whitney test (two-tailed) in (**c, d** and **l**). Difference between control and sni with stimulation were analyzed by unpaired t-test (**i, j**). ^***^ indicates *p* < 0.001, ^**^ indicates *p* < 0.01, ^*^ indicates *p* < 0.05 and n.s. means no significance.

Neuron types together making up the ensembles coding for mechanical and heat pain were next identified by scRNA-seq. As described above, mechanical and heat ensembles were captured in Fos^dsTVA*^R26^Tom^ mice by intraspinal delivery of EnvA^M21^ lenti-Cre virus co-incident with respective stimulation. Tom^+^ cells were sorted, scRNA-sequenced and mapped to the spinal cord atlas and plotted on top of the atlas (Fig. 3b) using label transfer^42^. Very few neurons from control group without stimulation were captured, sequenced and mapped to the reference atlas (Fig. 3b). Quantification of active neuron types in the different experimental conditions revealed an activation of the galanin (*Gal*) In8 neuron type as the only robustly activated inhibitory type and this was a shared feature for mechanical and heat pain (Fig. 3c). In contrast, among the activated excitatory neurons Ex21 and Ex19 were dominant for mechanical and heat stimuli, respectively (Fig. 3c). This result was independently validated using multiplexed RNA *in situ* hybridization of markers for these neuron types (Fig. 3d) revealing a co-expression of *Fos*;*Tac1* in mice subjected to mechanical stimulus, while co-expression of *Fos*;*Pou4f1* and *Fos*;*Tacr1* was found in mice subjected to heat stimulus (Fig. 3e and ExtendedData Fig. 6a, b). Although some deeper *Fos*^+^ neurons co-expressed these markers, most were lamina I neurons. Other neuron types (e.g. *Crhbp*^+^ Ex25 neurons, Fig. 3e) showed either no or proportionally small contributions to the ensembles. Spinoparabrachial (SPB) neurons that convey information related to mechanical and thermal noxious stimuli express *Tacr1, Gpr83* and/or *Tac1* and some of the neurons expressing these markers display unique supraspinal target innervation, even though *Tacr1*;*Gpr83* and *Tacr1*;*Tac1* are co-expressed in several neuronal populations^1,17-19,43^. *Tacr1*^+^ neurons are essential for responses to highly noxious stimuli and central sensitization^44,45^. We found that among the *Tacr1* expressing neuron types (Ex13, Ex17, In18, Ex19, Ex20) that *Gpr83* was expressed in Ex17 and Ex20, while expression of *Tac1* was detected on low level in several populations (In2, In3, Ex11, Ex17, Ex19, Ex20, Ex21 and Ex22) where Ex21 was marked by abundant expression (Fig. 3d). This is consistent with that *Tacr1*^+^;*Pou4f1*^+^;GPR83^-^ Ex19 represent SPB neurons coding for noxious heat while *Tac1*^+^ Ex21 SPB neurons code for intense mechanical stimuli (Fig. 3b-d). Beyond stimuli-selective activation of spinal neurons, several other excitatory and a few inhibitory neurons represented shared neurons in the ensembles (Fig. 3c). Thus, ensembles coding for noxious mechanical and heat stimuli included separate neural representations and predicted a single inhibitory molecular neuron type uniquely expressing *Gal* in the spinal cord to gate affective pain regardless of modality.

### Ensemble transition from nociceptive pain to chronic allodynia and hyperalgesia

To test whether the quality selective neural ensembles encoding mechanical and heat pain represent unique ensembles for nociceptive pain or whether neuropathic pain share these circuits, we scRNA-sequenced spinal cord segments corresponding the central arborizations of the sciatic nerve from mice subjected to spared nerve injury (sni) and control mice. The BAF53b-Cre^*^R26^Tom^ mouse strain was used to enrich for neurons. In total, 28,195 cells passed quality control of which 17,229 cells were identified as neurons and the rest were border associated macrophages (BAMs), microglia, oligodendrocytes and astrocytes (Fig. 4a). The sni animal model induces symptoms of neuropathic pain such as allodynia and hyperalgesia^46^. Neurons from both control and sni mice clustered into the same cell types as in the original reference neuronal atlas (Fig. 4b), with largely unaltered population size representation (ExtendedData Fig. 7a). However, differential gene expression analysis showed that up to thousand genes were affected in most neuronal types (Fig. 4c, Supplementary Tables 3 and 4). Machine learning based perturbation analysis^41^ further showed neuropathy to cause a marked effect on most neuronal cell types, whereas neurons from control mice were unaffected (ExtendedData Fig. 7b). Computational reconstruction of the gene regulatory network with SCENIC^47^ interestingly revealed broad effects on the activities of regulons (i.e. transcriptional regulator and cis-regulatory target genes) for several activity-induced genes, while other regulons were largely unaffected. The activity-dependent regulons (*Fos, Srf, Egr, Fosb, Junb, Jun*) were robustly increased in sni mice as compared to control animals in Ex10, E17, Ex19, Ex20, Ex22 and Ex25-E27 as well as In8 and In9 (Fig. 4d, ExtendedData Fig. 7c). *Fos* expression and *Fos* regulon module score analysed in the scRNA-sequencing data were increased in most if not all of these neuron types, a finding that was consistent with a marked increased expressing of *Fos* in ipsilateral spinal dorsal horn neurons as compared to contralateral to the lesion analysed by *in situ* hybridisation (Fig. 4e, f and ExtendedData Fig. 7d, e). This shows that neuropathy does not lead to subtle alterations in any selective neuron types, but instead a broad dis-inhibition of numerous excitatory neuronal types. This can only be explained by an altered environment, such as inflammation. We analysed microglia, macrophages, oligodendrocytes and astrocytes (ExtendedData Fig. 8 and 9). Cell clustering, differential gene expression and RNA velocity analyses on *Aif1*^*+*^ cells (microglia from control mice) showed microglia to mainly be resting (M0) or activated (M1, pro-inflammatory) and BAM (ExtendedData Fig. 8a-e). In the chronic phase after nerve injury, resting microglia were almost absent, and a proportional increase of activated microglia was observed (ExtendedData Fig. 8f-h). RNA velocity analysis revealed activated microglia to arise from the resting subtype (ExtendedData Fig. 8c). No subpopulation composition^48^ alterations were observed in oligodendrocytes (ExtendedData Fig. 9a-f), or astrocytes (ExtendedData Fig. 8g-i). Thus, microglia and macrophage activation could therefore explain the marked perturbation of spinal neurons during chronic stage of neuropathic pain, as has been proposed previously^49-53^. SCENIC analysis revealed increased activity of immediate early gene regulons in macrophages after sni and in activated M1 microglia compared to resting status (ExtendedData Fig. 8i). Further differential gene expression and gene ontology analysis confirmed a significant increase of antigen presentation via MHC class II and signalling for lymphocyte activation in macrophages (Extended-Data Fig. 8j, k) and enrichment of pro-inflammatory cytokines like *Ccl3, Ccl4* in resting (M0) and *Tnf, Il1b, Il6* in activated (M1) microglia in animals with neuropathic pain (ExtendedData Fig. 8l, m). This is consistent with a spinal inflammation driven by border associated macrophages and microglia during peripheral neuropathy. To examine functional consequences on chronic pain, neural ensembles representing mechanical and heat pain were first trapped for expression of hM4Di-DREADD followed by a peripheral nerve injury to induce neuropathic pain. Silencing the noxious mechanical and heat spinal nociceptive ensembles did not resolve nor relieve mechanical allodynia and hypersensitivity to blunt pricking (2g von Frey filament), heat and cold stimuli (Fig. 4g, h). These results indicate an inflammatory driven transition of ensembles from acute nociceptive pain to chronic allodynia and hyperalgesia. The transition manifests in a disinhibition of numerous spinal excitatory neuron types.

### Gal+ In8 neurons gates noxious mechanical and heat input and can reverse neuropathic pain

GABAergic interneurons are effectors of inhibitory modulation of primary afferent signalling. Loss of inhibitory input in the spinal dorsal horn has been shown to contribute to pain in physiological and pathological conditions^22,25,54^, and transplantation of GABAergic interneurons into spinal dorsal horn can ameliorate neuropathic pain^55^. *Gal*^+^ In8 neurons represented the major inhibitory neuron type in the mechanical and heat ensembles. Thus, the modality selective excitatory mechanical (Ex21) and heat (Ex19) neurons seemed both to be feed-forward gated by In8 neurons. We examined the impact of these neurons for nociception and neuropathic pain. *Gal* mRNA expression in wild type and *Cre*-dependent Tomato expression in Gal-Cre mice was mainly in neurons just superficial to/intermingled with *Prkcg* (PKCg) expressing neurons, thus in laminae I and II (Fig. 5a), consistent with previous spatial transcriptomics data of In8 neurons (Gaba2/3 in Häring’s annotation)^30^. The Gal-Cre driver mice faithfully recapitulated endogenous *Gal* expression, as there was a perfect co-localization of *Cre* and *Gal* mRNA expression in superficial layers of spinal cord (ExtendedData Fig. 10a). We delivered hM3D(Gq) AAV virus intraspinal in wild-type (Wt) and Gal-Cre mice (Fig. 5b) and examined coping and reflex behavior after administration of clozapine. Activation of *Gal*^+^ neurons in the lumbar spinal cord of Gal-Cre mice had no effect on heat withdrawal responses but caused an increased mechanical withdrawal threshold and decreased coping responses to both noxious mechanical and heat stimulations (Fig. 5c, d). While the effect was significant, the relative influence of the Gal-Cre neurons was greater in the more intense noxious hot plate than for the blunt pricking pain (2g von Frey filament).

We next sought to identify the afferent input to the Gal-Cre neurons. Retrograde rabies tracing from the spinal *Gal*^+^ neurons with AAV^TVA^ helper virus in Gal-Cre mice revealed monosynaptic input from a group of DRG neurons with medium-(44%, 300-700 μm^2^) to large-size (56%, >700 μm^2^) based on cross sectional area (Fig. 5e, f). We used several cell-type specific and broader makers for the different sensory neuron types^56-59^ to identify traced neurons (Fig. 5h). The traced neurons did not express markers for any of the unmyelinated nociceptive, thermoreceptive and pruriceptive sensory neurons. Most of the traced neurons were myelinated (*Nefh*^+^, 95%), and 38% were *Calca*^+^;*Gfra1*^+^, 27% were *Calca*^+^;*Gfra1*^-^ and 31% were *Calca*^*-*^;*Gfra1*^+^ (Fig. 5g, h and ExtendedData Fig. 10b). We classified the *Calca*^+^;*Gfra1*^-^ neurons as PEP2 and PEP3 neurons that are presumed A-heat sensitive and A-high threshold mechanoreceptive neurons^57^, respectively. The *Calca*^*-*^;*Gfra1*^+^ neurons were consistent with Ab-field LTMRs since traced neurons were negative for *Calb1* (marking Ab RA-LTMRs) and *Ntrk2* (marking Ad-LTMRs) but were *Slc17a7*^+^ which is exclusively expressed in touch sensitive Ab-LTMRs. Combined, this shows a monosynaptic input to Gal*-*Cre neurons from the A-fiber nociceptors PEP2, PEP3 and from Ab field-LTMRs.

Increasing the overall inhibitory tone in the spinal cord reverses neuropathic pain, suggesting that a reduced GABAergic tone may contribute to hyperalgesia symptoms^54,55^. Thus, it seemed possible that a decreased activity in In8 might explain disinhibition of excitatory neurons driving the pain pathway. Control mice and sni mice (18 days) received either a mechanical or a heat stimulus and were sacrificed 30 min later. Quantification of *Fos*^+^;*Gal*^+^ neurons in superficial laminae of the mice revealed a significant reduction of mechanical and heat recruitment of *Gal*^+^ neurons in mice with sni, as compared to control mice (Fig. 5i, j). This suggests that *Gal*^+^ neurons can contribute to the reduced spinal inhibitory tone and thus, restoring activity could reverse neuropathic pain. Mice with hM3D(Gq) expression in Gal-Cre spinal neurons (Fig. 5b) were subjected to the sni model of neuropathic pain (Fig. 5k). As compared to the basal threshold of behavioural response prior to sni (BT), sni animals had a marked increased coping behaviour to noxious mechanical, heat and cold stimuli, as well as reduced withdrawal thresholds to mechanical and heat (Fig. 5l). Activation of Gal-Cre neurons reversed neuropathy-induced pain-like behaviour in coping responses to mechanical, heat and cold, with smaller or no effect on withdrawal thresholds to mechanical and heat, respectively (Fig. 5l). Thus, despite inflammation in the spinal cord and the marked molecular perturbation of many populations of spinal neurons during sni, enhancing activity of Gal-Cre neurons was sufficient to reverse pain-like behaviour. The essential role of In8 in mouse made us examine whether this applies for cross species homolog neurons in the human spinal cord. Correlation analysis of gene regulatory networks (GRNs) between mouse and human spinal cord^60^ datasets revealed conserved *Gal*^+^ In8 population in human spinal cord (Inh-Dorsal-8 and Inh-Dorsal-6, ExtendedData Fig. 10c), suggesting that a similar gating mechanism could also exist in human.

## Discussion

Here, we examine the neural basis for one of the most fundamental mechanisms enabling a protection from harmful stimuli, the feeling of pain, by using several recent technological advances, including the high temporal resolution CANE approach for genetic trapping of neurons, scRNA-seq, machine learning classifiers, and functional manipulation of discreet ensembles and neuron types. We discover that neural activity alone within spinal ensembles is sufficient to recreate and completely abolish the sense of mechanical and heat pain. Thus, the identified ensembles are the carriers of cutaneous noxious mechanical and heat stimuli. Recent studies have revealed the involvement of molecularly different primary sensory neurons for the transduction of heat and mechanical stimuli. The affective component of noxious heat is probably carried by C- and A-fiber nociceptors expressing TRPV1. Withdrawal threshold to mechanical stimuli involves MrgprD^+^ C-fiber neurons, while the affective behaviour to noxious mechanical input also involves C-fiber TRPV1^+^ neurons as well as A-high-threshold mechanoreceptors (A-HTMR)^57^. While primary sensory neuron input involves several neuron types, the simplicity of spinal ensembles indicates these to encode activity among the primary sensory neuron types into discreet and dedicated circuits with only a few molecular neuron types responsible for several dimensions experienced by noxious external stimuli. Our results suggest that the cell-type composition of the ensembles represent stimuli modality, valence as well as intensity because subthreshold ensemble activation potentiated pain-related behaviour. Thus, intensity could be scaled by increasing active neuron numbers and/or firing frequency of already active neurons, as has previously been proposed^61^. However, the ensembles did not code for threshold detection and protective withdrawal responses, consistent with recent studies suggesting unique spinal neurons that code for reflex defensive reactions versus pain-related behaviour^17,62^. Our results show that mechanical and heat stimuli on the skin are represented by spinal ensembles with a weaker recruitment of excitatory neurons shared between stimuli and stronger recruitment of stimuli unique neuron types. Among the modality selective activated excitatory neurons, Ex21 defined by *Tac1* expression encodes intense noxious mechanical stimulus and Ex19 defined by *Pou4f1*;*Tacr1* expression represents intense heat. Spinal *Erbb4*^+^ interneurons have been recently shown to participate noxious heat sensation associated with protective reflexive responses, which governs heat sensation together with *Sst*^+^ and *Cck*^+^ neurons^16^. However, *Erbb4* is widely expressed in spinal cord such as In5, In9, Ex12-15, Ex17-19 and Ex27, which covers all excitatory neuronal populations together with *Sst* and *Cck* in current neuronal atlas. Mice with ablation of spinal *Tac1* expressing neurons display a loss of the affective behaviour associated with noxious stimuli, such as aversion and licking, but not protective reflexive responses^17^. This phenotype was observed after both noxious mechanical and heat stimuli suggesting that *Tac1*^+^ neurons have a general role in driving the affective component of sustained pain in a manner that is independent of sensory modality^17^. Therefore, our finding that the mechanical and heat ensembles control coping behaviour but not withdrawal threshold is consistent with the ablation of *Tac1* expressing spinal types. However, previous single cell RNA-sequencing has revealed *Tac1* expression at lower levels also in other populations of the dorsal spinal cord^30,32^, consistent with our new atlas (i.e. Ex17, Ex19-22). We find that ensembles including Ex19 and the Ex21 represent parallel pathways engaged by noxious heat and mechanical stimulation, respectively, and when chemogenetically re-activated, heat and mechanical-induced responses were facilitated without any cross-modality spill-over. Thus, even when activity is artificially forced, the ensembles stay tuned. This shows that the ensembles represent discrete processing units required and sufficient by themselves to code for noxious mechanical and heat stimuli. Therefore, we conclude mechanical and heat pain to be encoded by highly specialized local neuronal networks in the spinal cord. Within these ensembles, Ex19 and Ex21 may be projection neurons, as *Tacr1* and *Tac1* are present in spino-parabrachial neurons^1,17-19^.

Inhibitory neurons of the spinal dorsal horn exert critical control over the relay of mechanical and heat nociceptive signals to higher brain areas^22^. The spinal excitatory neuron types converge with feed-forward inhibition and the interdependence of the spinal network components together shapes the spinal output signal which carry information about environmental experiences conveyed by the primary sensory neurons^9,56^. While we found that heat and mechanical stimuli are represented by parallel processing units of excitatory neurons, both ensembles shared the In8 GABAergic neuron type. We interpret In8 as neurons gating excitatory output from the heat and mechanical ensembles because enhancing the activity of *Gal*^+^ In8 neurons ameliorated pain behavior to both mechanical and heat stimuli. In contrast, the lack of cross ensemble effects indicates that the mechanical and heat ensembles which both include In8 neurons, to be organized functional units, as has been shown for sensorimotor reflexes^62,63^. Consistently, indiscriminative activation of In8 blocked both mechanical and heat pain as well as injury-induced chronic pain. Our results contrasts to previously studied parvalbumin expressing dorsal horn interneurons (*Pvalb* expressed in In3, In4, Ex14, Ex15), and prodynorphin (*Pdyn*, expressed in In4 and In8) interneurons which have been shown to be involved in a circuit gating A-LTMR input involved in protective reflexes^14,25^ and directly inhibit spino-parabrachial neurons^64,65^. Thus, our results suggest that unlike parvalbumin and prodynorphin interneurons, *Gal*^+^ In8 represents a key inhibitory neuron type gating affective dimensions of pain with relatively marginal effects on protective reflexes. Likely because of this, the importance of *Gal*^+^ interneurons for nociception have gone undetected^66^. Inhibitory neurons expressing the potassium voltage-gated channel interacting protein 2 (*Kcnip2*) are activated preferentially by cold, but unlike *Gal*^+^ neurons have no impact on noxious mechanical or heat pain^67^. We speculate that spinal neurons gating reflex behavior were not TRAPed in this study because the protective reflex circuitry is activated already at the innocuous stimuli range and hence, activity-dependent expression of *Fos* may not be as robust as it is following intense noxious stimuli. The relation of function to other inhibitory interneurons previously functionally studied such as those expressing nitric oxide synthase 1, neuronal (*Nos1*)^66,68^ and neuropeptide Y (*Npy*)^69,70^ is difficult to assess as they are expressed in multiple excitatory and/or inhibitory neuron types. Our findings of direct monosynaptic connectivity with nociceptors are consistent with In8 as feed-forward inhibitory neurons in the pain pathway. Thus, presentation of a sensory stimulus recruits’ inhibition in addition to excitation, leaving an excitation/inhibition balance. Feed-forward inhibitory neurons in the spinal sensorimotor pathway are essential in the processing of touch stimuli and shapes the representation of touch in the somatosensory cortex^20,71^. Similarly, it is conceivable that In8 feed-forward inhibition improves acuity and contrast through lateral inhibition. Processing of pain-related information can be modulated by cognitive and emotional brain states^72^. Based on the impact of In8 neurons on output neurons and pain-related behaviours the neurons could also directly integrate supraspinal input onto the ascending pain pathway.

The nociceptive neurons can be categorized in terms of the preferred stimulus, threshold of activation and their conduction velocity. Heat sensation involves a range of cold, warmth and heat detecting skin receptors, with a dominance of slow C-over fast A-fiber heat nociceptors^73,74^. Similarly, high forces of mechanical stimuli result in the activation of a range of both touch and nociceptive neuronal types which are heat-insensitive, some of which are C-fibers and others A-fibers (A-high threshold mechanoreceptors, A-HTMR)^75,76^. Dorsal root ganglion neurons have been molecularly classified using single cell RNA-sequencing^56,58^. In the current taxonomy of mouse nociceptors and pruriceptors^57^, several molecularly defined C-fiber types have been shown to be heat or mechanical sensitive nociceptors while there is one predicted A-heat and one A-HTMR neuron type, named PEP2 and PEP3, respectively^57^. Interestingly, monosynaptic input to In8 is only through A-fiber nociceptors. Ab field-LTMRs are neurons that form large receptive fields terminating as circumferential endings around hair follicles with sensitivity to gentle skin stroking^77^. While C-fiber nociceptors are evolutionary conflated in human as compared to mouse (2 vs 4 types), A-fiber nociceptors have expanded and include four molecular types in humans while there are only two types in mouse^78^. Feed-forward inhibition of fast nociception might, thus, play a more important role in humans as compared to rodents.

Hyperalgesia caused by peripheral neuropathy involves a range of spinal excitatory neurons not normally conveying modality-selective mechanical and heat nociception. We draw this conclusion from analyses of active neurons during sni, the marked circuit-wide molecular perturbation, the failure of nociceptive ensembles functionally to respect noxious information quality and that inhibition of nociceptive ensembles unsuccessfully resolves allodynia and hypersensitivity in mice. Neuropathic pain resulting from peripheral nerve injury leads to colony-stimulating facto-1 (*Csf1*) released from primary afferents that drives microglia activation and a secondary neuroinflammation in the spinal cord^51^. Microglial cells have an important role in initiating and maintaining pain and inflammation^49,79-81^. Our results concur with these studies as we find a marked transition of resting state (M0) into activated (M1) microglia. We find increased expression of inflammatory cytokines in M1 microglia (including *Il1b, Tnf, Il6*) and downregulation of anti-inflammatory *Il10*. Interestingly, also M0 microglia displayed a range of gene expression changes including upregulation of the pro-inflammatory chemokines *Ccl3, Ccl4*, inflammation-induced transcription factor *Nr4a1* nuclear receptor which suppress expression of genes in the inflammatory NfkB pathway, as well as the NF-κB inhibitor-α (*Nfkbia*), which repress NF-κB and signalling^82,83^. Thus, there are molecular alterations of microglia which may contribute to the inflammatory state and hyperalgesia. BAMs (perivascular macrophages here) as professional antigen presenting cells together with activated microglia could orchestrate neuroinflammatory responses with infiltrated T lymphocytes to drive neuropathic pain after sni, even though meningeal macrophages (resolving pain) and infiltrated T cells (driving pain) have different roles in development of neuropathic pain^53,84^. The pro-inflammatory mediators are believed to enhance synaptic transmission to produce central sensitization and neuropathic pain^85,86^. Analysis of transcription factors and their putative targets unexpectedly did not reveal any gene-regulatory networks associated with inflammation in neurons. While these results do not exclude post-transcriptional alteration downstream of receptors activated by inflammatory mediators, it suggests that conventional inflammatory signalling pathways that are known to engage transcription are not responsible for gene-expression alterations in spinal neurons. Instead, we found activity-regulated gene transcription across many kinds of excitatory neurons, consistent with marked alterations in ensembles coding for nociception as compared to those active in animals with neuropathy. The activity-dependent gene transcription includes genes whose products are essential for synapse plasticity and long-lasting changes at synapses^87^. Such gene expression changes could beyond the microglial neuroinflammation contribute to maintenance of the hyperactive spinal circuits in the sni neuropathy model.

Several different kinds of inhibitory spinal cord dorsal horn neurons have been implicated in sensitization and mechanical allodynia in neuropathy including those that express prodynorphin^14^ and parvalbumin^25,88^. However, it is important to note that these markers are not exclusive to a single discreet inhibitory interneuron type, and several are expressed also in excitatory neurons as shown by neurochemical studies^89^ as well as single cell RNA-sequencing efforts, such as our current study. While *Gal*^+^ In8 neurons as part of the mechanical and heat nociceptive ensembles were unable to reverse neuropathy-induced hypersensitivity, forced activation of Gal-Cre neurons reversed mechanical, heat and cold pain-related behaviours and unlike prodynorphin and parvalbumin neurons, only marginally affected mechanical allodynia (and heat-induced protective reflexes). The failure of intense mechanical or heat stimulation to fully activate *Gal*^+^ neurons in sni mice as compared to control mice suggests that an In8-dependent break in the excitation/inhibition balance in sni drives a disinhibition of excitatory neurons. While Galanin receptors *Galr2* and *Galr3* are not expressed in spinal neurons, we find the inhibitory *Galr1* receptor conspicuously expressed specifically in Ex21, Ex23 and Ex24 populations. Galanin administered intrathecally attenuates neuropathic pain behaviour in rats^90-92^, hence, both Gal and GABA released from In8 neurons could bring about the excitatory/inhibitory balance.

Comprehensive strategies for pain relief across pain types are urgently needed^93,94^. Our results provide a cellular basis for how affective properties of acute pain perception caused by cutaneous noxious stimuli is encoded in the spinal cord. This could be helpful for development of therapeutic strategies targeting affective pain while preserving sensory processes for the detection of noxious stimuli and for protective reflexes.

## Materials and Methods

### Animals

Wild type female and male C57BL/6 mice (adult, ∼12 wk of age) were obtained from Charles River (Scanbur AB). Knock-in mice strains Fos^dsTVA^ (Fos-2A-dsTVA, strain #: 027831), Rosa26^Tom^ (Ai9, strain #: 007909), Rosa26^RC::FPDi^ (RC::FPDi, strain #: 029040) and Rosa26^DTA^ (strain #: 009669) were obtained from The Jackson Laboratory (JAX). Transgenic mice strains ACTB^FLPe^ (strain #: 003800, JAX, from Ole Kiehn), Gal-Cre (KI87, GENSAT, from Tomas Hökfelt), Actl6b-Cre (BAF53b-Cre, strain #: 027826, JAX, from Ulrika Marklund) were obtained at Karolinska Institutet locally. Rosa26^PDi^ mice (G_i/o_ protein-coupled receptor DREADD hM4Di driven by Cre) were obtained by crossing Rosa26^RC::FPDi^ and ACTB^FLPe^ mice. Fos^dsTVA*^R26^PDi^ mice (heterozygous for both gene loci) were obtained by crossing Fos^dsTVA^ and Rosa26^PDi^ homozygous mice. Fos^dsTVA*^R26^Tom^ mice (heterozygous for both gene loci) were obtained by crossing Fos^dsTVA^ and Rosa26^Tom^ homozygous mice. Fos^dsTVA*^R26^DTA^ mice (heterozygous for both gene loci) were obtained by crossing Fos^dsTVA^ and Rosa26^DTA^ homozygous mice. BAF53b-Cre^*^R26^Tom^ mice were obtained by crossing BAF53b-Cre and homozygous Rosa26^Tom^ mice. The mice were kept under standard conditions on a 12/12-h light-dark cycle with free access to food and water. The experiments were conducted in accordance with Swedish policy for the use of research animals and were approved by a local ethical committee (Stockholms Norra djurföröksetiska nämnd). Efforts were made to minimize the number of mice used and their suffering throughout experiments.

### Genotyping

DNA was extracted from a small piece of mouse ear tissue with sodium hydroxide and Tris^95^. Primers and programs used for genotyping were acquired from JAX genotyping protocols for strains from JAX. The primers and program used for Gal-Cre mouse strain were acquired from Mutant Mouse Resource & Research Centers (MMRRC).

### Viruses

Lenti-cre virus pseudotyped with mutated EnvA^M21^ (R213A; R223A; R224A, titer 4.7 ×10^9^ - 2.0 × 10^10^ vg(viral genomes)/ml) was produced by Duke viral vector core (Duke University) based on the protocol from Sakurai et al^33^. AAV9/2-hEF1a-DIO-hM3D(Gq)-mCherry (v98-9, titer 5.2 × 10^12^ vg/ml) viruses were produced by viral vector facility in University of Zürich. AAV9/2-Syn-flex-TVA.E66T-P2A-GFP-P2A-oG-WPRE3 (7.2 ×10^12^ vg/ml) and EnvA RV Δ G-mCherry (1.4 × 10^8^ IU (infection units)/ml) viruses were produced by viral vector facility in Charité University (pAAV-hSyn-FLEX-TVA-P2A-EGFP-2A-oG was a gift from Salk investigator John Naughton and BRVenvA-1n Rabies Virus, pseudotyped EnvA, mCherry was a gift from Edward Callaway (Addgene plasmid #32630, #32631, #32632, #32633,#32634))^96^.

### Peripheral sensory stimulations

Heat or mechanical stimulation to the left hind paw of mouse was performed under deep anesthesia with isoflurane (Baxter). For heat stimulation, the left hind paw of the mouse dipped into the water bath (51.0 - 51.5°C) for 15 sec for two times with an interval of 1 min. For mechanical stimulation, mechanical force (600 g) was applied to the hind paw by Randall Selitto Paw Pressure Meter (World Precision Instruments) for 10 sec and repeated 5-10 times with an interval of 1 min. Fos activation by sensory stimulations could be visualized at mRNA level (fresh tissue/30 min after stimulation) with RNAscope or protein level (perfused tissue/2 hr after stimulation) with immunohistochemistry. Intra-spinal injections of pseudotyped lenti-cre virus were applied for capturing activated ensembles after sensory stimulations. For comparison of the active ensembles captured by lenti-virus and Fos labelling, a second round of the same sensory stimulations were applied and the animals were allowed to survive for 2 hr before perfusion.

### Intra-spinal injections

Surgical procedures were performed under anesthesia with isoflurane according to previous description^22^ after 30 min of peripheral sensory stimulations. Briefly, an incision (2-3 cm long) was made on the back skin to expose T13 (thoracic) and L2 (lumbar) vertebrae, which corresponds to lumbar L4-L6 spinal cord segments). The mouse was then mounted to a motorized stereotaxic frame (Stoelting and Neurostar) and the vertebral column (L1) was fixed with spinal adaptors (Harvard Apparatus). Muscles, spinous process and left vertebral lamina were removed to expose one side of L5 segment (L4-L6). Small bleedings could be stopped with absorbable hemostatic gelatin sponge (Spongostan, Agnthos). The dura in the middle position of L5 segment was pierced about 400 µm left to the middle anterior spinal artery with a bevelled 30G needle (BD Microlance). Viral vectors were injected at a depth of 300 µm with a pulled glass micropipette mounted on a 10 µl Hamilton syringe. A 10 min injection (30 nl/min, 300 nl) process was controlled by an Elite nanomite pump (Pump 11, Harvard Apparatus). The glass micropipette was kept in place for 5 min after injection. Another two injections were made on rostral and caudal both directions about 1.2 mm away from the first injection site. Skin was closed with 5-0 silk stitches (Ethicon, Agntho’s). Xylocaine and Carprofen (Apoteket, Sweden) was used to relief pain caused by surgery. The mice were ready for experiments after 2-3 weeks. For monosynaptic retrograde tracing experiment with AAV helper virus and EnvA pseudotyped rabies virus, a slightly modified intra-spinal injection protocol was applied. A 1 cm incision was made through the skin to expose the gap between T13 and L1 vertebrae. The spinal column was fixed with a clamp (NARISHIGE, Japan) through the stabilization of L1 transverse process. The posterior longitudinal ligament and ligament flavum connecting T13 with L1 were cut to expose the spinal cord. AAV helper virus (500 nl) was injected at 400 µm right to the midline at a depth of 400 um on one side (one injection) and pseudotyped rabies (500 nl) was injected after 8 days of the injection of helper virus. The mice were allowed to survive for another 7 days.

### Spared nerve injury (sni)

Surgical procedures were performed under anesthesia with isoflurane as previously described^46,97^. Briefly, the skin of the mid-thigh from left lateral surface was incised, and a separation was made directly through the biceps femoris muscle exposing three terminal branches of the sciatic nerve: common peroneal, tibial and sural nerves. The common peroneal and tibial nerves were tightly ligated with one bundle of 6-0 silk (Ethicon), transected together distally to the ligation and a piece of 1-2 mm of the nerve was removed from the distal stump avoiding any contact with or stretching of the intact sural nerve during the surgery process. Finally, muscle and skin were closed in two layers with 5-0 silk stitches. Xylocaine and Carprofen was used to relief pain caused by surgery.

### Drugs

Clozapine (l mg/ml in ethanol as stock solution) was delivered through intraperitoneal (i.p.) injections (0.2 mg/kg in saline) for hM4D(Gi) mediated inhibition and hM3D(Gq) mediated excitation experiments. Behavioral tests were performed after 1 hr of administration of clozapine. For combined excitation experiment through hM3D(Gq) together with peripheral sensory tests, clozapine dosage was measured by titration, and the final dosage 0.04 mg/kg was applied. Combined behavioral tests were arranged after 2.5 hr administration of clozapine, where the effect of clozapine was considered as subthreshold.

### Single cell suspension preparation

Mice with a Cre dependent red fluorescent protein variant tdTomato reporter (10-20 weeks old, both males and females) were sacrificed with over dosage of isoflurane. Lumbar spinal cord was dissected out immediately and kept in freshly oxygenated artificial cerebrospinal fluid solution (modified ACSF: 87 mM NaCl, 2.5 mM KCl, 1.25 mM NaH_2_PO_4_, 26 mM NaHCO_3_, 75 mM Sucrose, 10 mM Glucose, 0.5 mM CaCl_2_, 4 mM MgSO_4_) on ice. Gray matter from lumbar spinal dorsal horn (or only lumbar 4-6 from ipsilateral side) was dissected out and transferred to a plastic petri dish (3 cm) with 2 mL pre-warmed digestion enzyme in ACSF solution (Papain, 25 unit/mL and DNase I, 55 unit/mL, Worthington Biochemical). For spinal dorsal horn neuronal atlas construction, each experiment included 2-3 mice (BAF53b-Cre^*^R26^Tom^). For labeled active ensembles or neurons from Sni mice, 4 mice were included for each suspension experiment. The following single cell suspension was performed according to our previous protocol^30^ with modifications. Briefly, spinal cord tissues were cut into pieces and triturated (10 × up and down) every 10 min (4 ×) using glass Pasteur pipettes with decreasing diameter (pre-coated with 0.5% BSA). Cell suspension was filtered through a 30 µm cell strainer (CellTrics, Sysmex) and washed with additional 1.5 mL ACSF and 0.5 mL PBS. Cells were spin down with centrifugation (300 g × 5 min, 4 °C) and resuspended with 2 mL ACSF. Cell suspension was carefully loaded on top of the same volume of OptiPrep density gradient medium (5.4% iodixanol in ACSF, Sigma) and centrifuge with 300 g × 10 min at 4 °C. Gradient centrifugation step was replaced with centrifugation (300 g × 5 min, 4 °C) for some samples. Cell pellet was resuspended with 4 mL ACSF. SYTOX Blue (Invitrogen, ThermoFisher Scientific) was added to stain dead cells. Then tdTomato positive and SYTOX Blue negative cells were sorted with fluorescence activated cell sorting (FACS) cell sorters (BD FACSAria Fusion / BD FACSAria III) at 4 °C. Cells were concentrated when necessary, by centrifugation (300 g × 5 min, 4 °C) and resuspended with proper volume (∼1,000 cells/ µl) of ACSF solution.

### Single cell gene expression 3’ sequencing

Sorted cells were loaded onto the 10 × Chromium chip G to yield single cell droplet with v3 or v3.1 kit (10 × genomics). Targeted cell recovery for neuronal atlas construction or Sni samples was settled to 5,000 cells, whereas the samples from captured active ensembles were aiming for 1,000 cells. Reverse transcription, cDNA amplification and library construction were performed according to the user guide provided by manufacture. Pooled libraries were sequenced on Illumina sequencing platform NovaSeq 6000 system on SP-100 flowcells with 91 bp sequenced into the 3’ end (5’ to 3’) of the mRNAs in National Genomics Infrastructure (SciLifeLab). Raw sequencing data were de-multiplexed, converted into fastq format, and aligned to mouse reference mm10 using the STAR aligner to generate the gene-cell matrices.

### Single cell RNA-sequencing data analysis

Gene expression matrices were imported into R (4.1.0) and analysed with Seurat (4.0.6) with standard pipelines (Satijalab). For constructing spinal neuronal atlas, individual cells were filtered out from the dataset if they had fewer than 2,000 genes or more than 20% ratio of mitochondrial genes. Raw counts were normalized by a global-scaling normalization method “LogNomalize” that normalizes the feature expression measurements for each cell by the total expression, multiplies this by a scale factor 10,000 and natural log-transforms (log1p) the results. Highly variable features were identified using FindVariableFeatures() function (2,000 features by default) for the following analysis. Counts were centered and scaled for each gene. The effects of total UMI and percent of mitochondrial genes in each cell were regressed out using a linear model in the Scaledata() function. The top 50 principal components (PCs) were retrieved with the RunPCA() function using default parameters. JackStraw() function and ElbowPlot() function were combined to determine the dimensionality of the dataset and for the following clustering. Clustering was done with FindClusters() function using the shared nearest neighbour (SNN) modularity optimization technique (Louvain algorithm by default). In order to avoid possible over clustering, we chose an approach where the cells were clustered from the highest level of separation through the adjustment of dimensionality and resolution and followed by merging of transcriptionally highly similar clusters or separating functionally hybrid clusters. Non-neuronal cells were filtered out according to the expression of neuronal mark genes *Rbfox3* and *Snap25*. Finally, 27 clusters have been produced for spinal neuronal atlas with reduction = “pca”, dims = 1:25, resolution = 0.5. The non-linear dimensional reduction technique UMAP was used to visualize cell clusters. Cluster specific marker genes were identified with FindAllMarkers() function. Wilcoxon Rank Sum test was selected to identify differentially genes with at least 0.25 increasing logfc.threshold. Specific gene markers including canonical and new were selected from the list of differentially expressed genes for unbiased classified cell clusters.

For non-neuronal cells from Sni and control samples, individual cells were filtered out from the dataset (only protein coding genes) if they had fewer than 1,000 genes, less than 4,000 UMIs, or more than 10% ratio of mitochondrial genes. The same pipeline described above was applied, where the dims (1:5) and resolution (0.4) were adjusted for oligodendrocyte and microglia.

### Cell identity classification/prediction

The classifier was built using scPred (v. 1.9.2)^98^, which was based on a low-dimensional representation of gene expression, with a mixture discriminant analysis (MDA) as the underlying model^99^. Both the reference and query data were normalized with the same normalization method. The query data was aligned onto the reference data and classified using the pre-trained models. All neurons with a maximum prediction score greater than or equal to 0.55 were assigned the label of the highest scoring cell type and the unassigned cells were filtered out.

### Pseudobulk differential expression analysis

DESeq2^100^ was implemented here to perform pseudobulk differential expression analysis on a specific cell type cluster (sni vs control) with standard workflow in R^101^. The raw counts and metadata were extracted from the dataset containing sni and control groups. Counts at single-cell level were aggregated at the sample level for each cluster and the corresponding metadata at the sample level was also generated. Quality control at the sample level was performed with principal component analysis and hierarchical clustering methods based on normalized and regularized log transformed counts. DESeq2 differential expression analysis was performed, and dispersion estimates were plotted. The results for contrasted groups (sni relative to control) were adjusted through shrinking the log2 fold changes using the apeglm method^102^. Results of significant genes were filtered with adjusted *p*-value < 0.01 and 50% increase or 50% decrease.

### SCENIC analysis

Single-cell regulatory network inference and clustering (SCENIC)^47^ was implemented here to explore the effect of sni on cell status for each cell cluster on RNA-seq data with standard pipelines (Aertslab). The input data for SCENIC was a single-cell RNA-seq expression matrix containing 11,990 cells for neuronal clusters (5,995 cells per condition), whereas the expression matrix for microglia/macrophages remained the original cell numbers (1,368 cells for control and 969 cells for sni). Co-expression network was inferred by running GENIE3. Gene regulatory network (GRN, regulon) was built based on co-expression modules and transcription factor motif analysis (RcisTarget). Top50perTarget and top3sd were applied for CoexMethod while creating regulons for neuronal and glial datasets, respectively. Regulons were scored in the cells with AUCell. Only non-extended regulons from neuronal dataset were kept for regulon activity visualization. Fos module score was produced with the list of genes in Fos regulon through AddModuleScore() in R.

### Perturbation analysis

To identify cell types most affected by sni, we performed a perturbation using the R package Augur^103^. First, raw scRNA-seq data from SNI animals was processed and the cell labels were assigned from the reference data using scPred as described above. After this, the reference and SNI datasets were merged, and the full dataset was filtered to include only protein coding genes while also excluding any genes located on the Y-chromosome. The data were then integrated with Seurat to remove any batch effects between samples. Finally, Augur was run between control and SNI datasets for each cell type using default settings. As a negative control, we ran another round of Augur analysis on the same merged dataset with randomly shuffled treatment labels.

### RNA velocity

The M25 version (GRCm38.p6) from GENCODE was used as reference for mouse sequencing data. According to the reference sequence, we generated the composite kallisto index of the separate fragments for spliced and unspliced transcripts. The spliced and unspliced counts of each cell were quantified following the kallisto bustools workflow for 10× scRNAseq^104^. Cells were selected with the criteria of 2,000 expressed genes in each cell and at least 20 counts (both unspliced and spliced) for a gene. The over-dispersed genes were normalized, and log transformed. The total PCAs were selected as 30, and data was calculated via balanced KNN imputation as 30. Next, the dynamical model from scVelo^105^ was applied for estimating the velocities. The embedding scatter plot was forked from the UMAP plot generated by Seurat analysis. For better visualization, the scaling factor in Gaussian kernel around grid point was set as 0.35, and the threshold for mass to be exhibited were set to above two.

### Cross-species correlation analysis of gene regulatory networks (GRNs)

The cross-species correlation analysis of GNRs was conducted using the scCAMEL toolkit as described previously^78^. Briefly, the scCAMEL toolkit (https://sccamel.readthedocs.io/) was utilized for estimating cell type similarity and integrating data across species. The SWAPLINE package within scCAMEL calculates probabilistic scores for cell types by training a neural network model on the most differentially expressed genes, excluding cell cycle-related genes. These genes are normalized and used for model training, with accuracy validated through k-fold cross-validation. For cross-species data integration, scCAMEL employs interpretable neural-network learning. Each dataset is sequentially used as a reference for cell type prediction in other datasets. All the training-prediction results are merged for principal component analysis to identify significant components, which are then used for illustrating cell-type similarities. Additionally, gene expression normalization aligns gene symbols across species, and transcription factor-related gene regulatory networks are identified by using GENIE3 and facilitating further correlation analysis.

### Behavioral tests

Animals were habituated to the testing environment twice before behavioral experiments started. Mechanical withdrawal threshold was tested in clear glass (8 cm in diameter) on a metal mesh floor and measured by a logarithmically incremental stiffness of 0.04, 0.07, 0.16, 0.40, 0.60, 1.0, and 2.0 (g) von Frey Filament (Stoelting) combined with an up–down method to assess tactile allodynia^46,106,107^. The cut-off of a 2.0 hair was selected as the upper limit for testing. Heat sensitivity was assessed with Plantar Test (Hargreaves Method, IITC Inc. Life Science), and the response latency was recorded^108^. Cold plantar test was performed with a dry ice pellet (9 mm, AGA AB) under the glass (same settings of Hargreaves test), and the response latency was recorded^109^. Hot plate test was assessed with temperature at 50 °C (or 46 °C, specified in text) and the coping response latency (paw shaking, lifting/guarding or licking) was recorded.

To quantitatively scale pain responses, measures of coping episodes, durations or numbers of coping responses (paw shaking, lifting/guarding or licking) were applied. Mechanical stimulated response was tested with a 2.0 g von Frey filament for three times (successfully applied), and the average duration of coping behaviours was calculated. Alternatively, mechanical stimulated response was tested with a 1.0 g von Frey filament or 2.0 g von Frey filament, and the duration of coping behaviours was recorded. Mechanical hyperalgesia was also tested with a safety pin (23G needle, BD), and the duration of coping behaviors was recorded ^46^. Response to cooling stimulation was tested with a drop of acetone, and the duration of coping behaviors was recorded ^46^. Cold stimulated response was tested with a drop of pre-cold acetone on dry ice, and the duration of coping behaviors was recorded ^46^. Heat stimulated response was tested by hot plate, and the duration of coping behaviours was recorded for 3 min (46 °C) or 60 sec (50 °C). Sni induced heat hypersensitivity was tested by a water drop (50 °C) applied to the lateral hindpaw of animals on mesh floor through a 20 ml syringe.

### Tissue preparation and immunohistochemistry

Mice were deeply anesthetized with sodium pentobarbital (50 mg/kg administered i.p.; Apoteket) and perfused transcardially with 4% paraformaldehyde freshly made from powder as previously described^110^. The lumbar (L4-L6) spinal cord was dissected out and postfixed in the same fixative for 90 min at 4 °C, followed by rinsing in 10% (wt/vol) sucrose in 0.1 M phosphate buffer containing 0.01% sodium azide (Merck) and 0.02% bacitracin (Sigma). The tissues were kept in 10% sucrose solution for 2 d at 4 °C. All trimmed tissues were embedded with optimal cutting temperature cryomount (HistoLab AB), frozen with liquid carbon dioxide through sublimation, and sectioned on a cryostat (NX70, Thermo Fisher Scientific) at a thickness of 20 µm for spinal cord. The sections were mounted onto Superfrost Plus microscope slides (VWR International) and stored at −20 °C until use.

Spinal cord sections were dried at RT for at least 30 min. Then the slides were pre-treated with xylene 2 times for 10 min each to remain tissue morphology to the greatest extend for antigen retrieval, rehydrate with downgraded ethanol, rinse with deionized water and PBS. Antigen retrieval treatment (95°C for 20 min) was added to expose masked epitopes with Target Retrieval Solution (Agilent Dako). After an hour blocking with 10% serum at RT, rabbit anti-Fos primary antibody (1:1000, Santa Cruz) was added and incubated in a humid chamber at 4°C for two days. Immunoreactivities were visualized using the TSA Plus Fluorescein kit (PerkinElmer) as previously described^110^. For double labelling, slides with TSA labelling of Fos were selected and rinsed in PBS for 20 min and then incubated with primary antibody against PKCγ (1:200, Santa Cruz) or TVA (1:200, from Andrew Leavitt) over 48 hr at 4 °C. The slides were first washed in PBS for 30 min and then incubated with donkey anti-rabbit IgG (H+L) highly cross-adsorbed secondary antibody Alexa Fluor™ 555 or 647 (1:800; Invitrogen, Thermo Fisher Scientific) at RT for 2 hr; after rinsing in PBS for 30 min, counterstaining with 4’,6-diamidino-2-phenylindole (DAPI, Thermo Fisher Scientific) or propidium iodide (PI, Sigma) was followed. Finally, slides were mounted with fluorescence mounting medium (Agilent Dako), dry overnight at RT and stored at −20 °C until imaging.

### Tissue preparation and in situ hybridization (RNAscope)

Mice were deeply anesthetized isoflurane and decapitated. Lumbar spinal cord and/or DRGs was dissected out, snap frozen on dry ice and kept at −80°C. Frozen tissues were thawed on ice briefly, trimmed, embedded with optimal cutting temperature cryomount, frozen with liquid carbon dioxide, and sectioned on a cryostat at a thickness of 20 µm for spinal cord and 12 µm for DRGs. The sections were mounted onto Superfrost Plus microscope slides and stored at −80 °C before use.

RNAscope assay was performed according to the protocol provided with RNAscope Multiplex Fluorescent Detection Kit v2 (Cat. No. 323110, ACDBio) with minor modifications, where the hydrogen peroxide treatment was skipped, protease III was used instead of protease IV and counterstaining with DAPI (1.0 µg/ml, 10 min at RT) was performed as mentioned above for immunohistochemistry. Slides were mounted after rinse in PBS with mounting medium (Agilent Dako), dried overnight at RT and stored at −20 °C until imaging. The probes included in this study were designed and provided by ACDBio as listed in Supplementary Table 5.

### Microscopy and image processing

Representative images were acquired from one airy unit pinhole on an LSM700 confocal laser-scanning microscope (Zeiss) equipped with EC Plan-Neofluar objectives with a magnification of 10 × and 20 × and N.A. of 0.30, and a water objective with a magnification of 40 × and N.A. of 1.40. Emission spectra for each dye were limited as follows: DAPI (<480 nm), Alexa Fluor 488/Flurorescein (505–540 nm), Alexa Fluor 555/PI/Cyanine 3 (560–610 nm), and Alexa Fluor 647/Cyanine 5 (>640 nm). For projection images, orthogonal z-stacks were acquired with a depth interval of 1 µm with an objective with a magnification of 20 × or 40 ×. Images were processed using ZEN2012 software (Zeiss) and Fiji (ImageJ, NIH). Quantification was done with Fiji. Multi-panel figures were assembled using Adobe Photoshop and Adobe Illustrator (Adobe Systems).

### Statistics

Behavior data of coping episodes or numbers was presented as median with interquartile range and assessed by two-tailed Mann-Whitney test. For other behavioral tests and quantification for immunohistochemistry or RNAscope, data were expressed as mean ± SD and assessed by ordinary one-way ANOVA followed by Bonferroni’s multiple comparisons post hoc analysis by using Prism 10 software (GraphPad). The criterion for statistical significance was P < 0.05.

## Acknowledgements

We thank Prof. Fan Wang (MIT, USA) for providing us the CANE technology and generous support to get it to work, Drs. Jana Sontheimer and Roland Baumgartner for supporting work in the lab, Dr. Hannah Weman for technical input regarding rabies tracing, Drs. Lijuan Hu and Michael Fatt for 10 × technical support and animal staff from Department of Comparative Medicine helping with animals (KM-B) at Karolinska Institutet. This research was funded by the European Research Council (DescendPain 101053091 to PE), the Swedish Medical Research Council (2019-00761 to PE and 2022-00960 to MCL), Brain foundation grants to MCL and Knut and Alice Wallenberg Scholar and project grants to PE. Fellowships from Brain foundation to MDZ and JS, Swedish Society for Medical Research (SSMF) to MDZ, JS and YH, Sigrid Jusélius Foundation to JK. We also acknowledge the National Genomics Infrastructure in Stockholm funded by Science for Life Laboratory and computations and data handling by the Swedish National Infrastructure for Computing (SNIC) at UPPMAX funded partially by the Swedish Research Council (2018-0597), the Biomedicum Imaging Core (BIC) facility and Biomedicum Flow Cytometry Core facility at Karolinska Institutet.

