## Supplemental File for "Neural ensembles that encode affective mechanical and heat pain in mouse spinal cord"

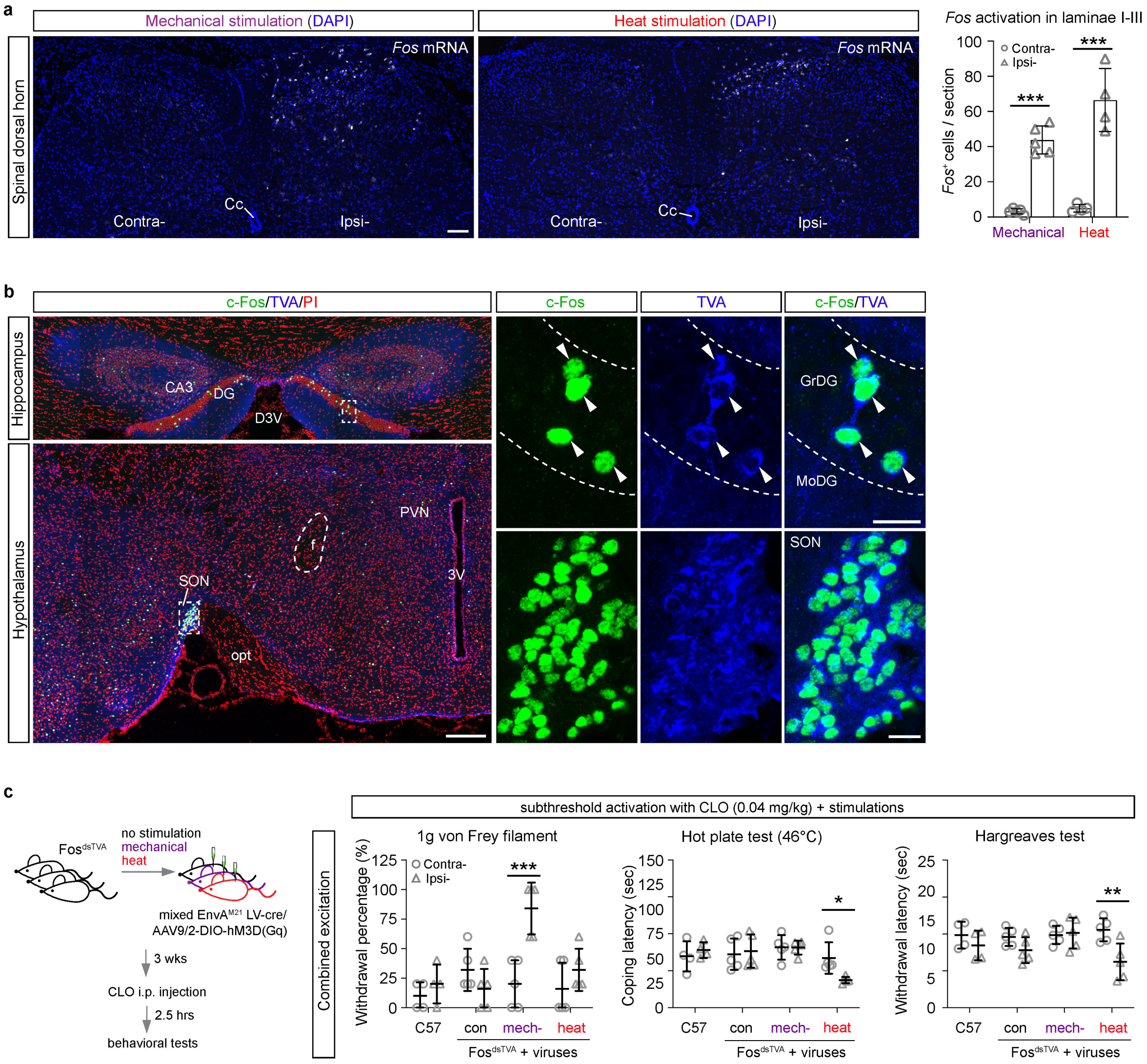


**ExtendedData Fig. 1: *Fos* activation by mechanical and heat stimuli in spinal dorsal horn, specificity of Fos$^{dsTA}$ mice and modulation of active neurons labelled by *Fos* expression**. **a**, Representative images of *Fos* mRNA expression in ipsilateral spinal dorsal horn induced by mechanical or heat stimulus. DAPI was used as a counterstaining for nuclei. Quantification of *Fos*+ cells (neurons) per section in laminae I-III was presented (n = 5 mice for mechanical and n = 4 mice for heat). Contra-, contralateral; Ipsi-, ipsilateral; Cc, central canal. **b**, The representative overview images of co-expression of c-Fos and TVA in hippocampus and hypothalamus from Fos$^{dsTVA}$ mouse brain counterstained with propidium iodide (PI). High magnification images show co-localization in granular cells of dentate gyrus (DG) and cells in supraoptic nucleus (SON) from the insets. GrDG, granule cell layer of the dentate gyrus; MoDG, molecular layer of the dentate gyrus; D3V, dorsal 3rd ventricle; opt, optic tract; PVN, paraventricular nucleus of hypothalamus; 3V, 3rd ventricle. Arrowheads indicate the granule cells express both c-Fos and TVA. **c**, Left, schematic diagram for the functional study of gain of function for ensembles infected by AAV9/2-DIO-hM3D(Gq) virus through subthreshold clozapine (CLO) combined with peripheral applied stimuli. Right, withdrawal percentage for 1g von Frey filament, coping latency for hot plate test (46°C) or withdrawal latency for cold plantar assay under subthreshold activation (0.04 mg/kg clozapine) of mechanical or heat ensembles (n = 19 mice). Con: Fos$^{dsTVA}$ mice injected virus without stimulation; C57: wild type mice; Contra-: contralateral; Ipsi-: ipsilateral. Scale bars = 100 μm in (A), 200 μm for the overview and 20 μm for high magnification in (**b**). Data are expressed as mean ± standard deviation. Differences between contra- and ipsi- from different groups were analyzed by ordinary one-way ANOVA followed by Bonferroni's multiple comparisons test (**a**, **c**). *** indicates $p < 0.001$, ** indicates $p < 0.01$ and * indicates $p < 0.05$.

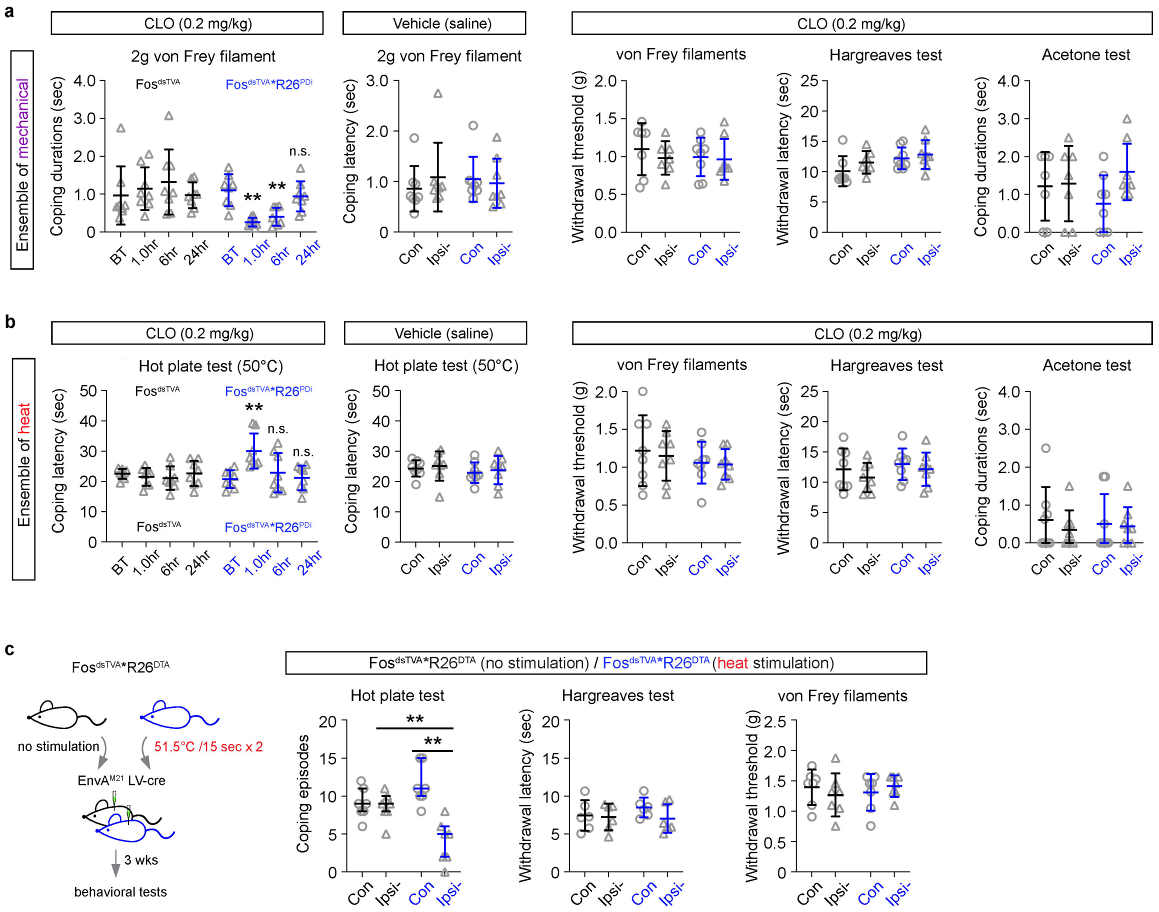


**ExtendedData Fig. 2: Effects of inhibition of ensembles coding for mechanical or heat pain in spinal cord with chemogenetics or loss of ensembles for heat with R26$^{DTA}$ mice.** Ensembles coding for mechanical or heat pain in spinal dorsal horn were captured with EnvA$^{M21}$ lenti-cre virus in Fos$^{dsTVA}$*R26$^{PDi}$ mice (**a** and **b**). **a**, Coping durations for mechanical stimulation was assessed with 2g von Frey filament at different time points (1.0 hr, 6 hr and 24 hr) after application of clozapine (CLO, 0.2 mg/kg, i.p.). Saline was used as the vehicle for clozapine (1.0 hr after injection). Withdrawal threshold for von Frey filaments test (non-noxious mechanical), Hargreaves test (thermo test) and coping durations for acetone test (non-noxious cold/cool) were assessed after 60 min of application of clozapine (n = 16 mice, 8 mice/group). Some tests only included 7 control mice. Fos$^{dsTVA}$ mice were used as a control group. BT, basal threshold. Con, contralateral. Ipsi-, ipsilateral. **b**, Coping latency for heat stimulation was assessed with hot plate test (heat, 50°C) at different time points (1.0 hr, 6 hr and 24 hr) after application of clozapine (0.2 mg/kg, i.p.). Saline was used as the vehicle for clozapine (1.0 hr after injection). Withdrawal threshold for von Frey filaments test, Hargreaves test and coping durations for acetone test were assessed after 60 min of application of clozapine (n = 16 mice, 8 mice/group). **c**, Left: schematic diagram for loss of spinal ensembles coding for heat pain on Fos$^{dsTVA}$*R26$^{DTA}$ mice. Right: coping episodes from hot plate test (50°C), withdrawal latency from Hargreaves test and mechanical threshold from von Frey filaments test were assessed and compared between contralateral and ipsilateral sides from heat stimulated and non-stimulated control groups (n = 14 mice, 7 mice/group). Hargreaves test only used 6 mice from each group. Data are expressed as mean ± standard deviation. Differences for different time points from ipsilateral and differences between contra- and ipsi- within the group were analyzed by ordinary one-way ANOVA followed by Bonferroni's multiple comparisons test (**a**-**c**). The coping episodes data in (**c**) are expressed as median with interquartile range and the differences between contra- and ipsi- from each group or ipsi- sides from different groups were analyzed by Mann-Whitney test (two-tailed). *** indicates $p < 0.001$, ** indicates $p < 0.01$ and n.s. means no significance.

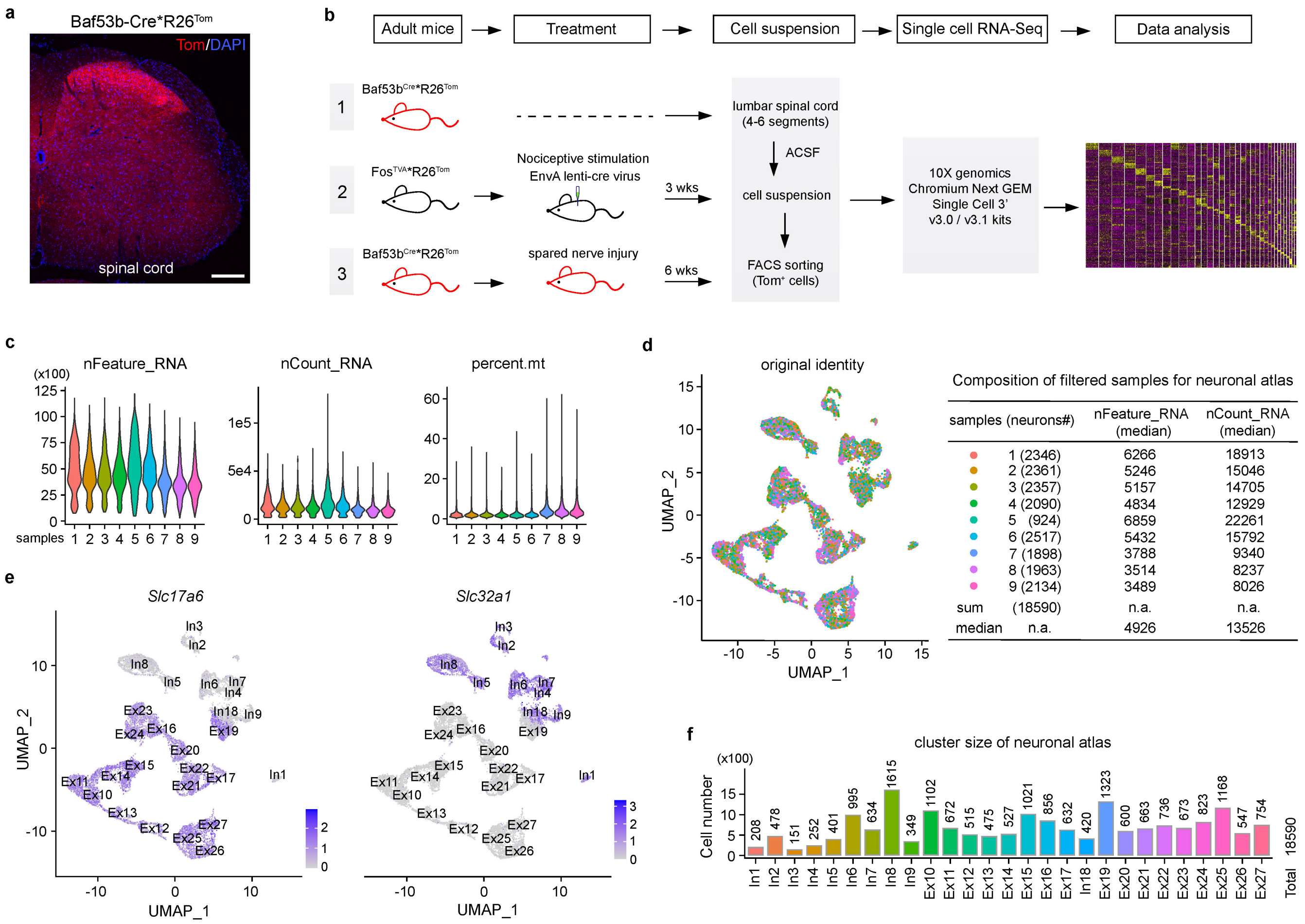


**ExtendedData Fig. 3: Workflow for single cell RNA-sequencing of neurons from adult mouse spinal cord and quality control of spinal neuronal atlas.** **a**, Representative image of Tom expression in spinal cord from Baf53b-Cre*R26Tom mice. DAPI was used as a counterstaining for nuclei. Scale bar = 200 μm. **b**, Experimental workflow of single cell suspension preparations from different scenarios and RNA-Seq included in this study. **c**, Overview of sequencing quality including number of detected genes (nFeature_RNA), total reads of detected genes (nCount_RNA) and percentage of mitochondrial genes (percent.mt) for 9 samples (without filtering) from adult mice spinal cord. **d**, Left: UMAP of spinal neuronal atlas composed of cells from different samples. Right: table for numbers of neurons contributed from each sample to spinal dorsal horn neuronal atlas and their corresponding median for number of detected genes and total reads of detected genes. **e**, UMAP plots of excitatory *Slc17a6* (vGLUT2) and inhibitory *Slc32a1* (VGAT) marker gene expression. **f**, Size distribution (number of cells) of individual cluster across the atlas, which was composed of 18,590 neurons.

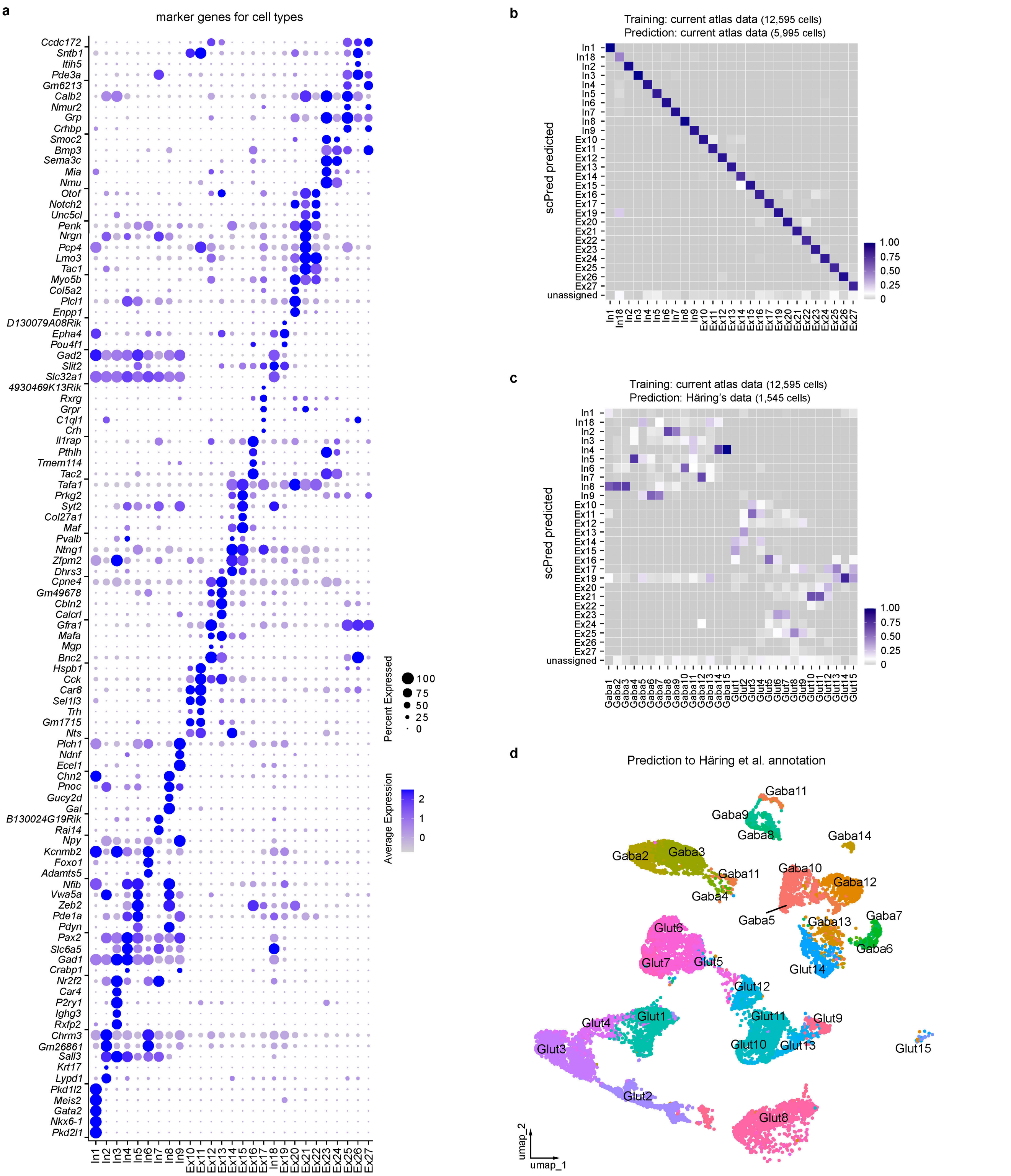


**ExtendedData Fig. 4: Expression of cluster marker genes of spinal dorsal horn neuronal atlas and comparison with Häring's annotation.** **a**, Dot Plot of top 5 genes for each cluster of the atlas. **b**, Heatmap of prediction of neurons from current atlas dataset with scPred to their known identity in the atlas. **c**, Heatmap of prediction of Häring's dataset to current atlas annotation with scPred. **d**, UMAP of neurons from current atlas with predicted Häring's annotation by Seurat label transfer.

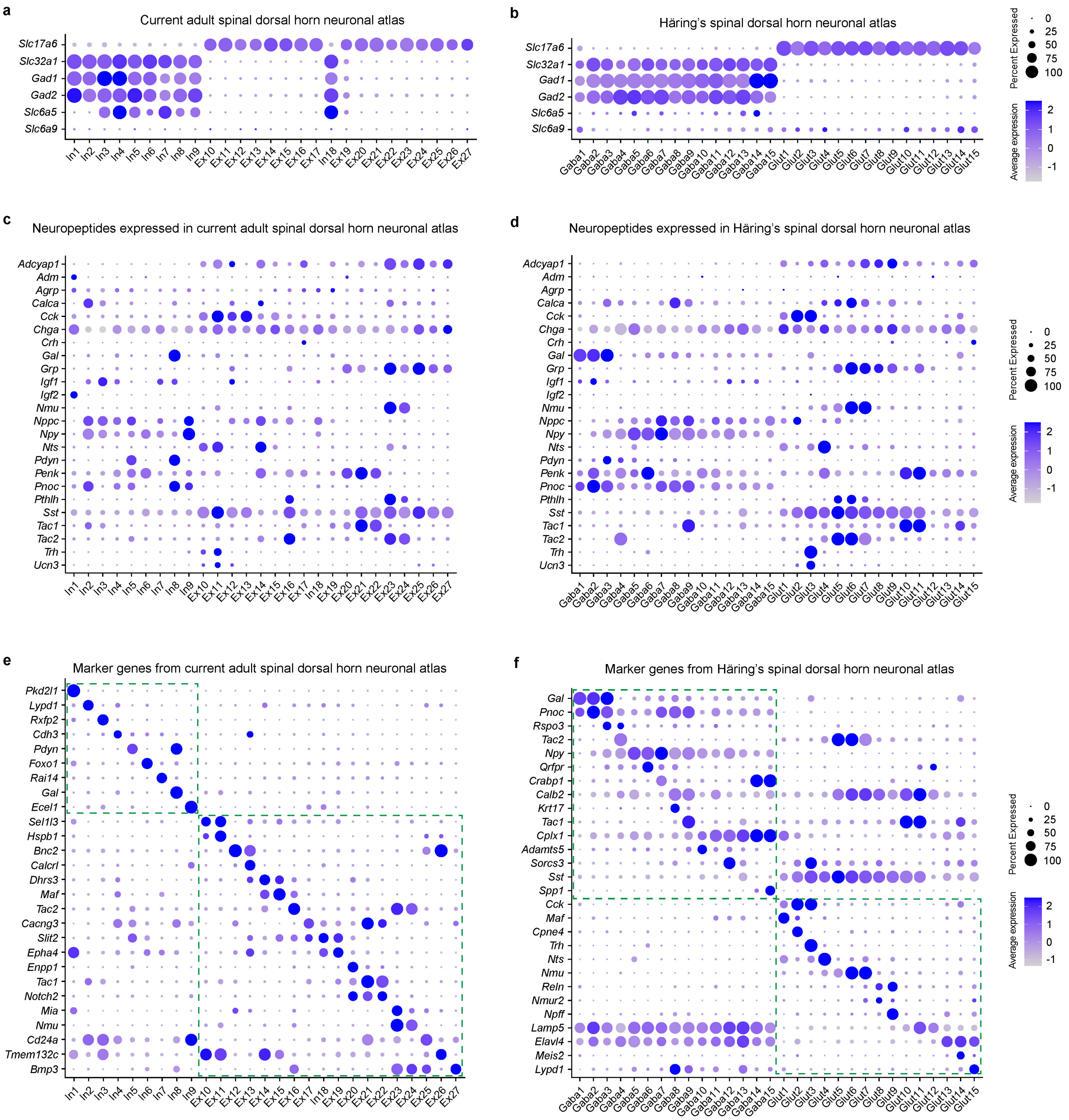


**ExtendedData Fig. 5: Comparison of expressions for excitatory/inhibitory genes, neuropeptides and lists of selected genes in current atlas and Häring's dataset.** **a** and **b**, Dot plots show expression of *Slc17a6*, *Slc32a1*, *Gad1* (GAD67), *Gad2* (GAD65), *Slc6a5* (GlyT2, neurons) and *Slc6a9* (GlyT1, predominantly astrocytes). **c** and **d**, Dot plots show expression of a list of neuropeptides. **e**, Dot plot shows selected marker genes for neuronal clusters of current atlas. **f**, Dot plot shows published marker genes in Häring's dataset. Dashed line rectangles highlight the enriched gene expression for inhibitory and excitatory clusters in (**e** and **f**).

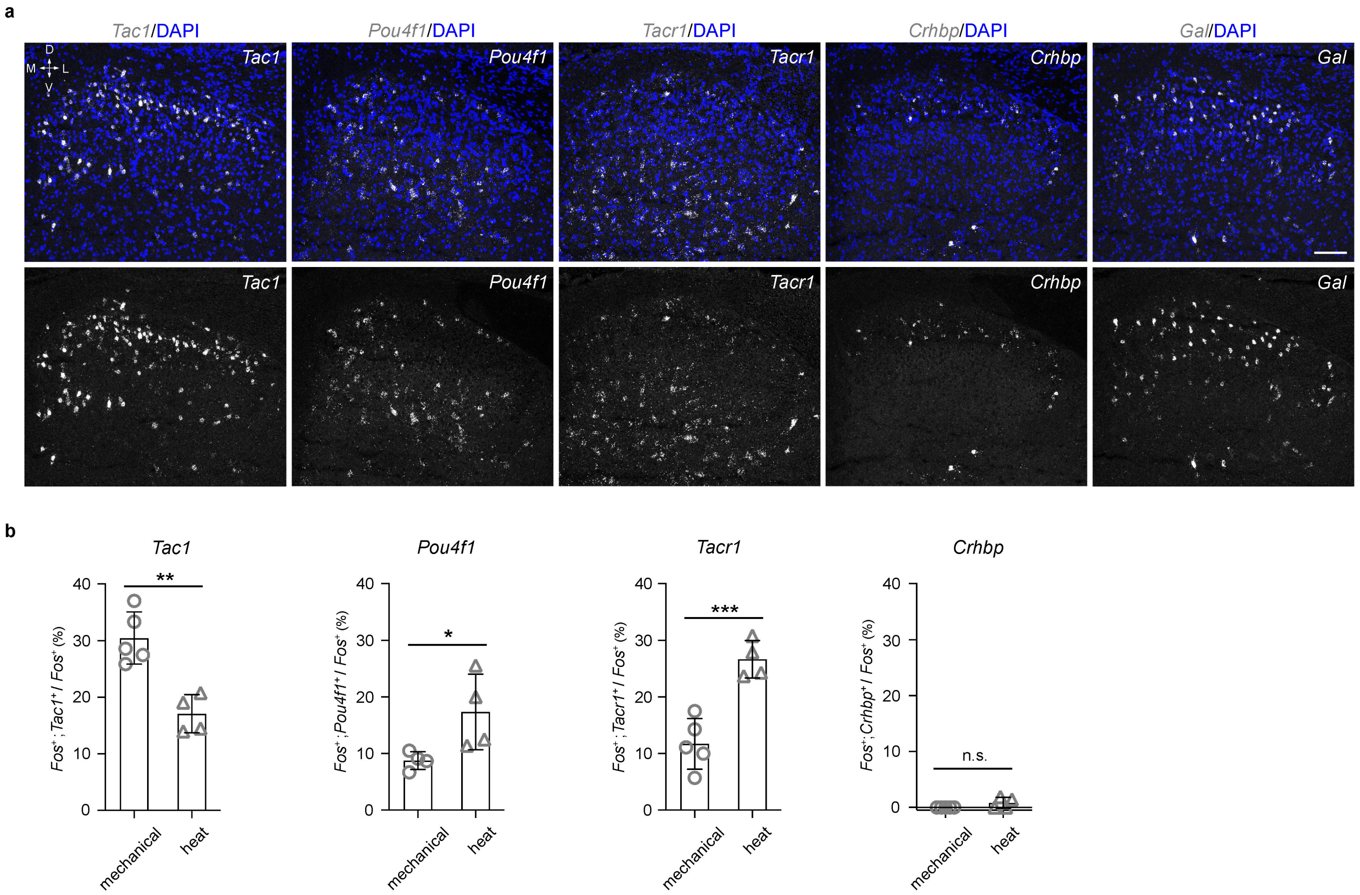


**ExtendedData Fig. 6: Expression of marker genes in mouse spinal dorsal horn and their co-localization with *Fos* stimulated by mechanical or heat. a**, Representative images of *Tac1*, *Pou4f1*, *Tacr1*, *Crhbp* and *Gal* expression in mouse lumbar 4/5 spinal dorsal horn (n = 4 mice). DAPI was used as a counterstaining for nuclei. Scale bar = 100 μm. M, medial; L, lateral; D, dorsal; V, ventral. **b**, Quantification of the proportion of co-localized *Tac1*, *Pou4f1*, *Tacr1* or *Crhbp* with *Fos* to $Fos^{+}$ neurons of spinal dorsal horn with mechanical or heat stimulation (n = 8 mice, 4 mice/group). Data are expressed as mean ± standard deviation. Differences between the stimuli were analyzed by unpaired t test (two-tailed). *** indicates $p < 0.001$, ** indicates $p < 0.01$, * indicates $p < 0.05$, and n.s. means no significance.

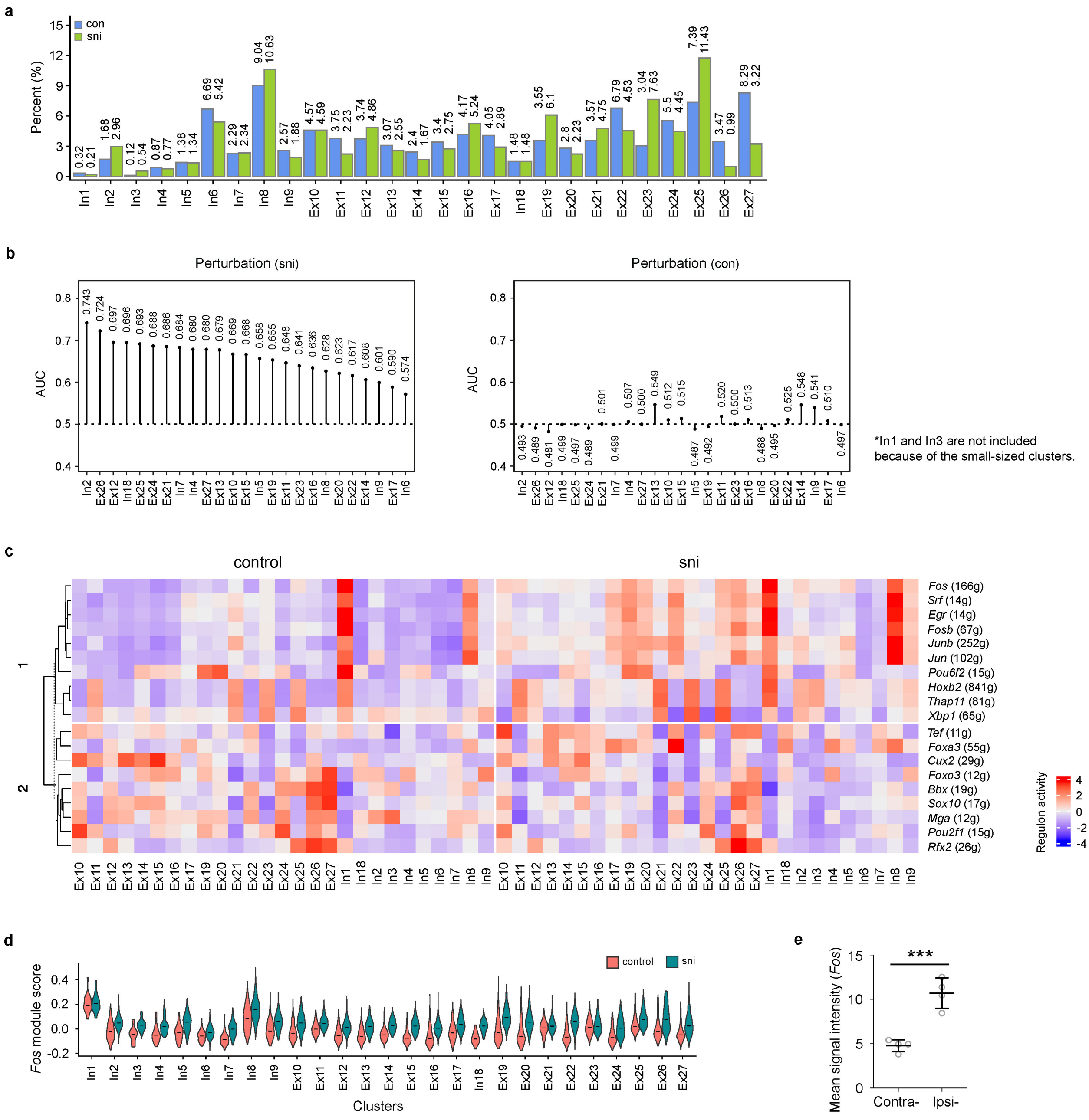

**ExtendedData Fig. 7: Effects of sni on neuronal cell types in mouse spinal dorsal horn. a**, Effect of sni on the size (proportion) of neuronal cell types in mouse spinal dorsal horn. Con, control. **b**, Left: Perturbation analysis of sni on neuronal subtypes in spinal dorsal horn with Augur method. Right: Perturbation analysis on neuronal subtypes from a randomly selected control sample as negative control was also included. **c**, Enriched regulons for spinal dorsal horn neuronal cell types from control and sni mice by SCENIC analysis. **d**, Violin plot of module score for the list of genes from *Fos* regulon (166 g) in neuronal cell types from control and sni. **e**, Quantification of mean signal intensity for *Fos* mRNA (subtracted background, n = 4 mice). Data are expressed as mean ± standard deviation. Differences between the stimuli were analyzed by unpaired t test (two-tailed). *** indicates $p < 0.001$.

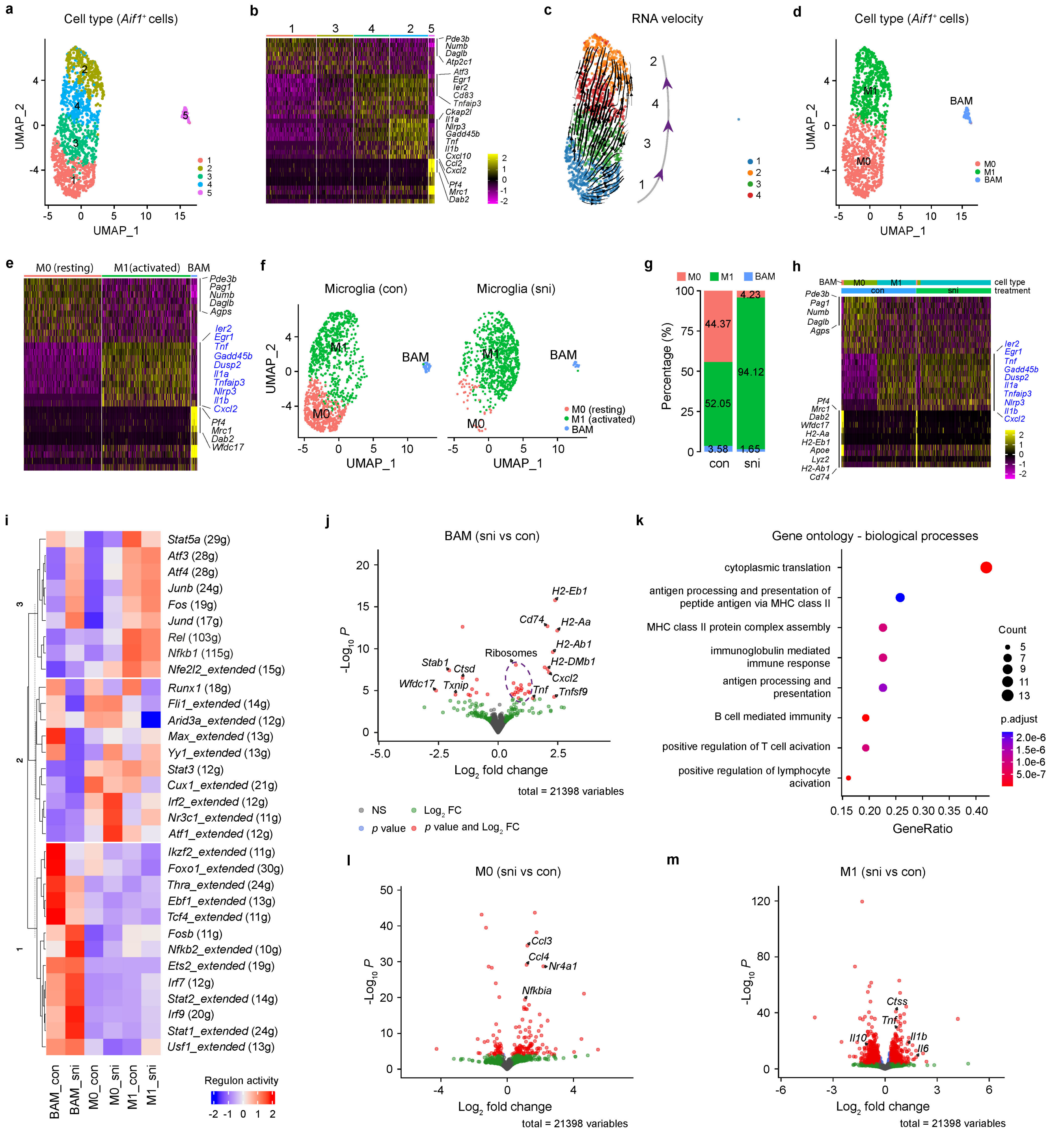


**ExtendedData Fig. 8: Effects of sni on *Aif1*+ cells in mouse spinal dorsal horn.** **a**, UMAP plot shows 5 clusters of *Aif1*+ cells (1,368 cells) from spinal cord of control mice. **b**, Heatmap shows top 10 differently expressed genes (avg_logFC) for each cluster. **c**, RNA velocity shows microglia activation tracjectory. Although cells of cluster 5 are *Aif1*+, they are border associated macrophages (BAM) according to specifically expressed genes in (**b**). **d**, UMAP plot shows the annotation of *Aif1*+ glia cells (M0, resting microglia; M1, activated microglia) according to top differently expressed genes, RNA velocity analysis and literature. **e**, Heatmap shows top 10 differently expressed genes enriched based on the newly annotated glial subtypes (M0, M1 and BAM). **f**, UMAP plots show the composition of glial cells from control and sni mice. **g**, Stacked bar plot shows proportion of different glial cell types from control (1,368 cells) and sni (969 cells) mice. **h**, Heatmap shows top 10 differently expressed genes from BAM, M0 and M1 cell types under control and sni conditions. Only 950 glial cells from each sample were included for visualization by down sampling. **i**, Heatmap of regulon activity enriched for glial cell types. **j**, Volcano plot shows sni induced regulated genes BAMs. Dashed line circle indicates genes coding for ribosomal proteins. **k**, Gene ontology analysis revealed sni regulated genes in BAMs were mainly involved in protein translation and immune response processes. **l**, Volcano plot shows sni induced regulated genes in resting microglia (M0). **m**, Volcano plot shows sni induced regulated genes in activated microglia (M1). Pro-inflammatory cytokine genes *Ccl3*, *Ccl4*, *Il1b* and *Il6* or anti-inflammatory cytokine gene *Il10* were indicated with arrows in (**l** and **m**).

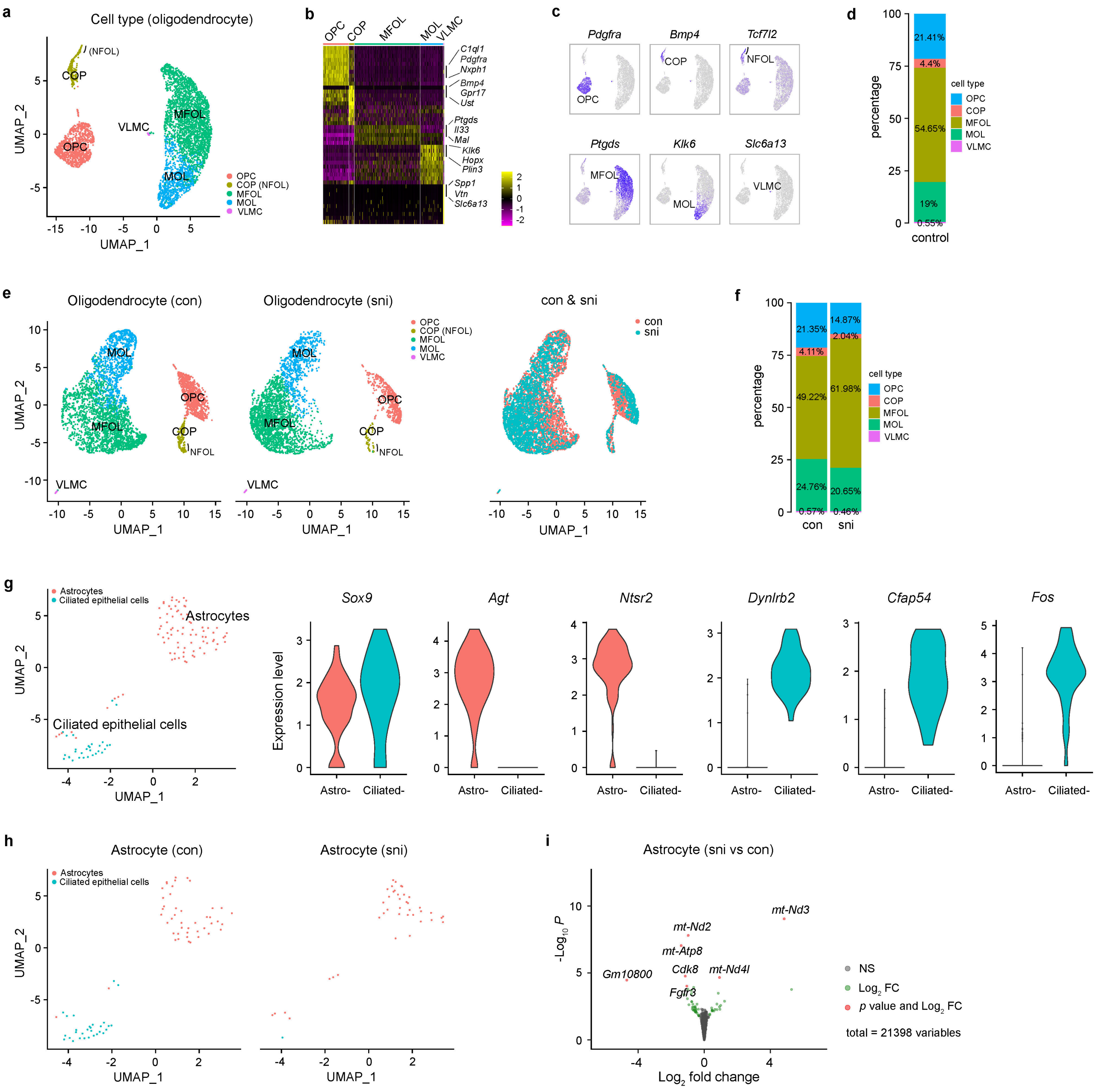


**ExtendedData Fig. 9: Effects of sni on oligodendrocytes and astrocytes in mouse spinal dorsal horn. a**, UMAP plot shows different cell subtypes along oligodendrocyte lineage from mouse spinal cord based on selected markers of Marques's 2016 annotation. OPC, oligodendrocyte precursor cells; COP, differentiation-committed oligodendrocyte precursors; NFOL, newly formed oligodendrocytes; MFOL, myelin-forming oligodendrocytes; MOL, mature oligodendrocytes. **b**, Heatmap shows top 10 differently expressed genes (avg_logFC) for different oligodendrocyte subtypes. **c**, Feature plots show selected marker genes for different oligodendrocyte subtypes. **d**, Stacked bar plot shows the population sizes of different oligodendrocyte subtypes. **e**, UMAP plots show oligodendrocytes subtypes from spinal cord of control mice (4,750 cells) and mice with sni (3,719 cells). **f**, Stacked bar plot shows the population sizes of different oligodendrocyte subtypes from con and sni conditions. **g**, Left: UMAP plot shows 2 clusters of *Sox9*+ cells (116 cells) from spinal cord of control and sni mice. Right: Violin plots show *Agt* and *Ntsr2* are specifically expressed in astrocytes, whereas *Dynlrb2* and *Cfap54* are limited to ciliated epithelial cells (ependymal cells). *Fos* is highly expressed in ciliated epithelial cells, like CSF contacting neurons (In1, **Fig. 4e**). **h**, UMAP plots show astrocytes and ciliated epithelial cells from spinal cord of control mice (74 cells) and mice with sni (32 cells). **i**, Volcano plot shows sni induced gene expression in astrocytes.

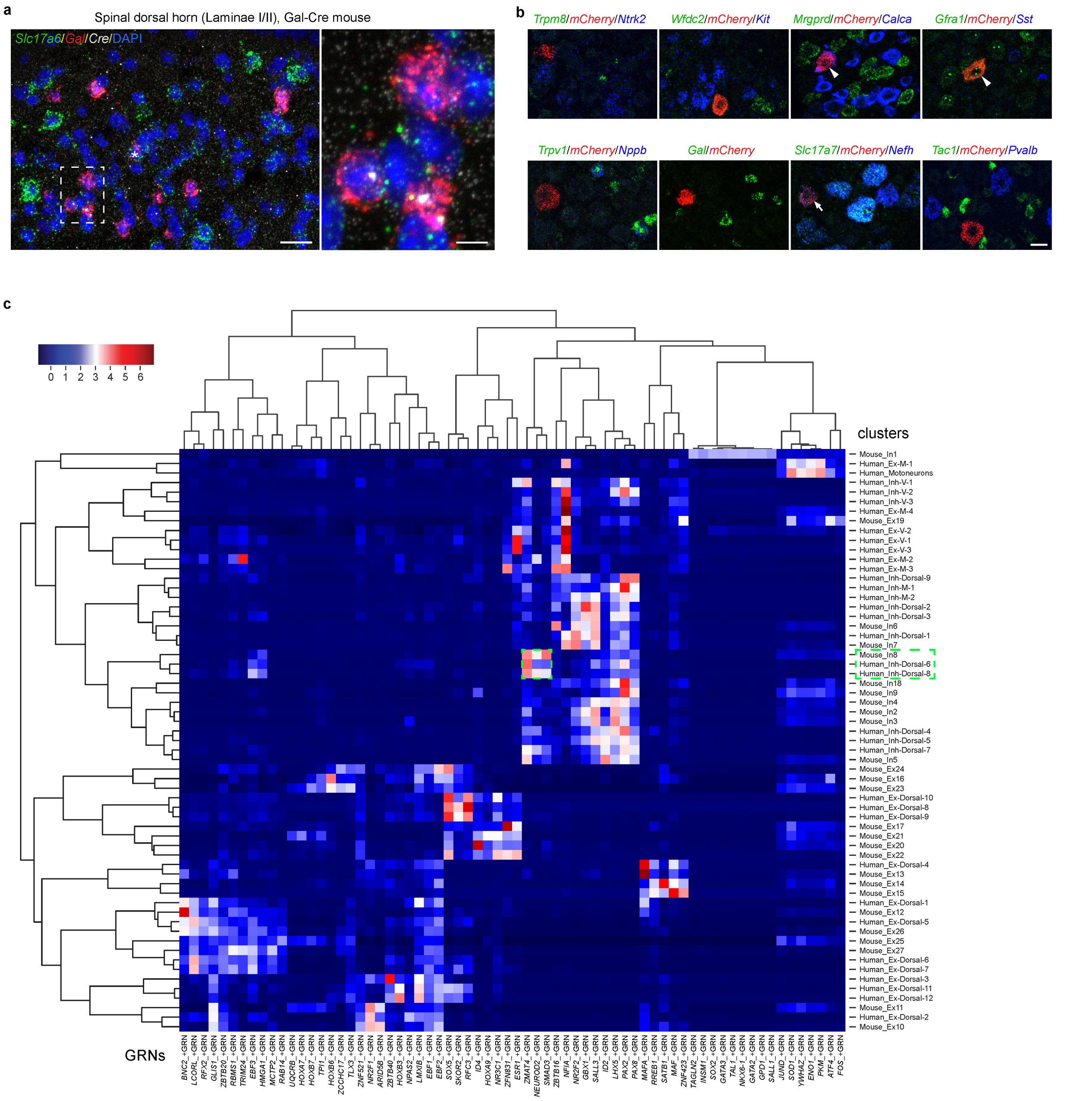


**ExtendedData Fig. 10: Identification of retrograde traced DRG neurons projecting to *Gal*+ In8 neurons and comparison between mouse and human spinal cord. a**, Representative image for the expression of *Slc17a6*, *Gal* and *Cre* in spinal dorsal horn from Gal-Cre mice (n = 3 mice). High magnification images show co-localization of *Gal* and *Cre* from the rectangle inset. Asterisk indicated the triple labeled (*Slc17a6*+/*Gal*+/*Cre*+) neuron. **b**, Identification of traced mCherry+ DRG neurons with combinations of RNAscope probes covering DRG neuronal subtypes. Arrows indicate triple labelling and arrowheads indicate double labelling. **c**, Heatmap with hierarchical clustering shows similar gene regulatory networks (GRNs) between mouse In8 and human Inh-Dorsal-6/8 according to Yadav's 2023 annotation. Scale bars = 20 µm in (**a**), 5 µm for the inset from (**a**) and 25 µm in (**b**).
